# Leveraging BRG1 Driven Ferroptosis Resistance to Overcome Treatment Resistance

**DOI:** 10.1101/2025.09.02.673601

**Authors:** Soo-Yeon Hwang, Helgi Nikolli, Belem Yoval-Sánchez, Sieun Yang, Peter Martin, Caitlin Gribbin, Lalit Sehgal, Lapo Alinari, Robert A. Baiocchi, Inah Hwang, Xiangao Huang, Maurizio DiLiberto, Alexander Galkin, Hojoong Kwak, Selina Chen-Kiang, Hongwu Zheng, Jihye Paik

**Author notes:** Correspondence: Jihye Paik, Ph.D, Associate Professor of Pathology and Laboratory Medicine, Hongwu Zheng, Ph.D, Assistant Professor of Pathology and Laboratory Medicine, Weill Cornell Medicine 1300 York Avenue, Box 69 New York, NY 10021.

## Abstract

Resistance to Bruton’s tyrosine kinase inhibitors (BTKi) remains a major therapeutic challenge in B-cell malignancies, limiting treatment durability. Here, we identify ferroptosis suppression as a central mechanism of BTKi resistance in mantle cell lymphoma (MCL). Aberrant BRG1 activity protects cells from BTKi-induced ferroptosis by restricting reactive oxygen species (ROS) and labile iron. Mechanistically, BRG1 promotes resistance through both BTK-dependent survival signaling and a BTK-independent transcriptional program. The latter is mediated by BRG1-driven induction of MEF2B, which upregulates NDUFA4L2 to inhibit mitochondrial respiration, thereby blocking mitochondria-dependent ferroptosis. Pharmacologic inhibition of BRG1 disrupts these programs, restoring ferroptotic sensitivity and synergizing with BTKi across resistant MCL models. Together, these findings establish BRG1 as a central regulator of therapy resistance and provide a rationale for co-targeting BRG1 and BTK as a therapeutic strategy for B-cell malignancies.

## Introduction

B-cell malignancies are life-threatening diseases with substantial unmet clinical needs ^1^. Bruton tyrosine kinase inhibitors (BTKi), first approved a decade ago, revolutionized the therapy by targeting a key intermediary in the B cell receptor (BCR) signaling pathway. BTKi have significantly improved outcomes in chronic lymphocytic leukemia, mantle cell lymphoma (MCL), Waldenström’s macroglobulinemia, and marginal zone lymphoma ^2–5^. In MCL, BTKi are now integral to frontline treatment and are increasingly associated with improved overall survival ^6–8^. However, despite high initial response rates, resistance develops in most patients within 1–2 years, leading to poor outcomes and a median survival of 2.9 month after ibrutinib (IB) failure ^6^. As BTKi move earlier into treatment course, the number of patients with BTKi-refractory MCL is expected to rise. Defining the molecular mechanisms of BTKi resistance is therefore critical for developing strategies to overcome treatment failure.

Resistance to BTKi in MCL arises from diverse mechanisms including genetic lesions (e.g. *BTK*, *BIRC3*, *TRAF2/3,* or *CARD11* mutations), adaptive pathway reactivation (e.g. NF-κB, PI3K/AKT/mTORC1, or MAPK), metabolic rewiring toward oxidative phosphorylation or tumor microenvironment-derived cues (e.g. cytokine signals and integrin-mediated stromal adhesion) that compensate for the BTK inhibition ^9^. To overcome the resistance, various strategies have been explored, including next-generation BTKi, and targeted agents against PI3K ^10–12^, BCL2 ^13–15^, and CDK4/6 ^16–21^, used alone or with BTKi. However, most have shown only modest, short-lived clinical benefits or treatment-limiting toxicities ^22,23^. Among these, BCL2 inhibitor has achieved ∼75% overall response rate in refractory MCL ^13,14^. However, many patients remain unresponsive, suggesting resistance mechanisms beyond evading apoptosis, potentially involving suppression of other cell death pathways.

BTKi resistance in MCL is rarely explained by genetic alterations in BTK itself, suggesting alternative mechanisms such as epigenetic reprogramming. Recurrent mutations in chromatin regulators, altered chromatin landscapes, and emerging dependencies on specific transcriptional networks ^24^, point to epigenetic dysregulation as a central driver of therapeutic resistance. BRG1 (encoded by *SMARCA4*), is a catalytic subunit of the mammalian BAF (SWI/SNF) nucleosome remodeling complex that regulates gene expression programs controlling proliferation, differentiation, DNA repair, stress responses, and therapy resistance ^25–29^. Its genetic alterations occur in ∼5-7% of human cancers ^30,31^, exhibiting context-dependent roles as either a tumor suppressor or oncogene ^32–35^. In B-cell lymphomas, BRG1 frequently exhibits missense mutations or haplo-insufficiency, while complete deletion is rare ^36,37^. In Burkitt lymphoma, its heterozygous missense mutations were found in 27% of cases ^38^. In genetic studies using mice, BRG1 haploinsufficiency drives lymphomagenesis from germinal center centrocytes, by disrupting transcriptional activity of SPI1, NF-κB, and IRF family, promoting a hyperproliferative state through BCL6 and MYC activation ^36^. In MCL, BRG1 forms a complex with SOX11 to activate oncogenic transcriptional programs that directly regulate the expression of key genes involved in disease pathogenesis ^39^. BRG1 mutations are enriched in BTKi-refractory MCL, present in 50% of IB-venetoclax resistant cases ^14,40^. BRG1 deficiency promotes resistance by derepressing anti-apoptotic programs, such as BCL-xL, via ATF3 silencing ^40^. While these findings reveal BRG1-mediated vulnerabilities, it remains unclear how BRG1-dependent transcription program promotes BTKi resistance.

In this study, we delineate a BRG1-mediated mechanism of BTKi resistance in MCL rooted in ferroptosis suppression. We uncover that cells expressing cancer-associated BRG1 mutants escape BTKi-induced ferroptosis by restricting intracellular labile iron and reactive oxygen species (ROS). This protection is mediated through both BTK-dependent signaling and a BRG1–MEF2B–NDUFA4L2 pathway that blocks mitochondria-dependent ferroptosis. Functionally, BRG1 sustains BCR survival signaling and MEF2B expression, while MEF2B-driven induction of NDUFA4L2 impairs mitochondrial activity to promote ferroptosis resistance. Importantly, we demonstrate that pharmacologic BRG1 inhibition restores ferroptotic sensitivity and synergizes with BTKi across resistant MCL models, establishing BRG1 as a central regulator of BTKi resistance and a promising therapeutic target.

## Results

### Ferroptosis suppression underlies BTK inhibitor resistance

To investigate the mechanism underlying BTKi treatment-induced MCL cell death, MINO and JEKO1 cells were co-treated with BTKi and the inhibitors of individual cell death pathways. Notably, co-treatment of the synthetic ferroptosis inhibitor, ferrostatin-1 (Fer-1) significantly suppressed BTKi-induced cell death (IB/PB/ZB *vs.* IB/PB/ZB+Fer1; 73.3/58.4/41.2% *vs.* 6.7/6.5/5.8% in MINO; 56.5/44.9/89.4% vs. 4.4/8.3/13.2% in JEKO1) (Fig. 1A, S1A). By comparison, the pan-caspase inhibitor Q-VD-OPh produced only a minor effect (Fig. 1A). These results indicate ferroptosis as the major pathway underlying BTKi-induced MCL cell death. Consistently, BTKi treatment markedly enhanced intracellular labile Fe²⁺ levels, a conducive condition to ferroptosis (Fig. 1B). The treated cells also exhibited elevated total reactive oxygen species (ROS) levels (Fig. 1C) and significantly increased lipid peroxidation (up to ∼7-fold in MINO and ∼10-fold in JEKO1) (Fig. 1D), indicating accumulation of iron-dependent oxidative damage, a hallmark of ferroptotic cell death. This was further supported by treatment of antioxidants β-mercaptoethanol or N-acetylcysteine which markedly suppressed IB-induced cell death (Fig. 1E). Finally, CRISPR-mediated BTK depletion also significantly elevated cellular levels of labile Fe²⁺ (Fig. S1B), total ROS (Fig. S1C), and lipid peroxidation (Fig. S1D), further demonstrating a protective role of BTK activity against ferroptosis.

**Fig. 1.**
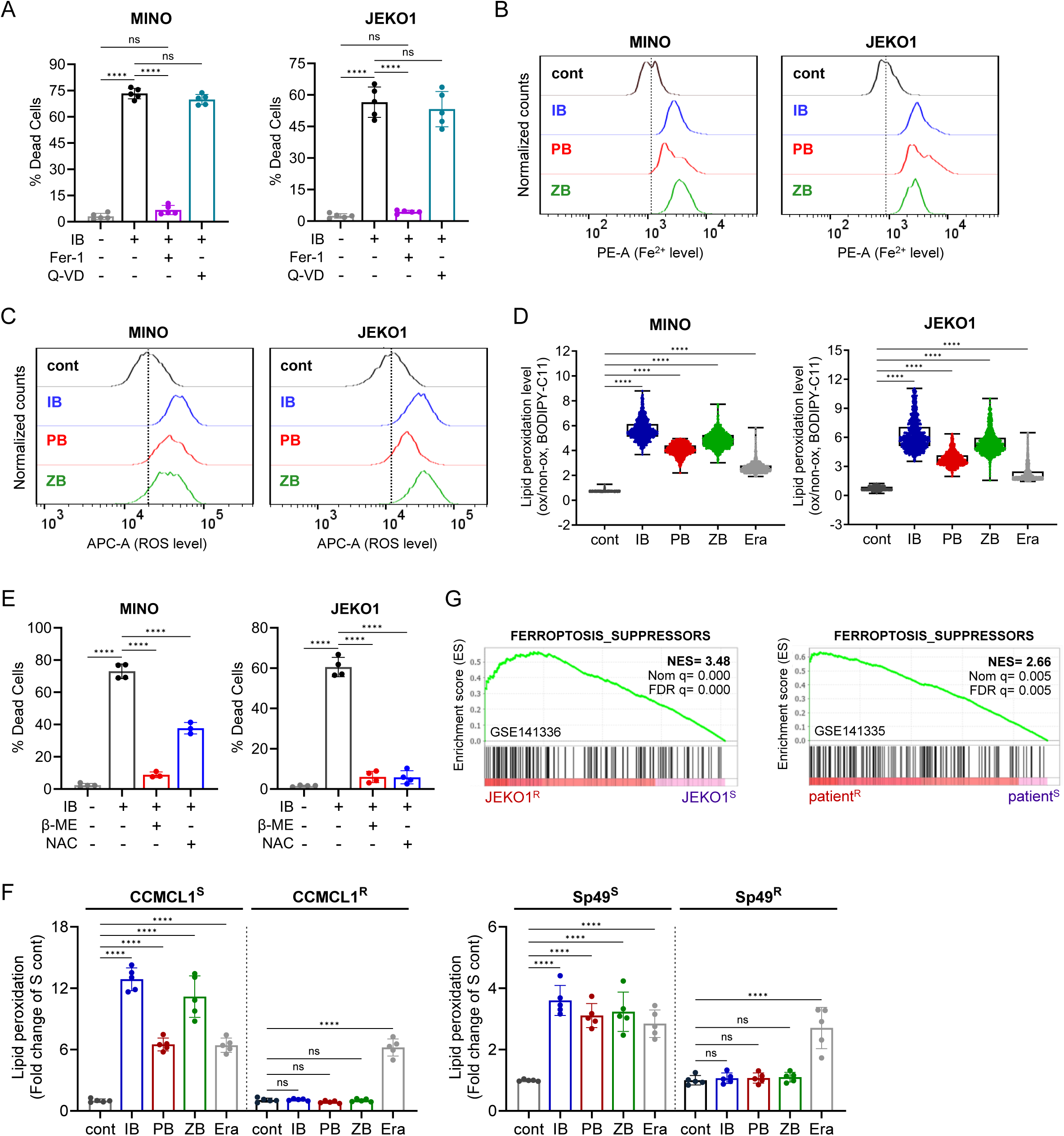
Ferroptosis suppression underlies BTK inhibitor resistance. (A) Effect of cell death inhibitors (Fer-1, 2 μM; Q-VD-Oph, 10 μM) on IB-induced (96 h) cell death in MINO and JEKO1 cells. (B-C) Representative flow cytometry histograms showing changes in labile Fe^2+^ levels (B) and ROS (C) following BTKi treatment (48 h). (D) BTKi-induced changes in lipid peroxidation (72 h) in BRG1^WT^ MCL cells. Erastin (0.5 μM) was used as positive control. (E) Effect of antioxidants (β-mercaptoethanol and N-Acetylcysteine; 100 μM) against IB-induced (96 h) cell death in BRG1^WT^ MCL cells. (F) BTKi-induced changes in lipid peroxidation (24 h) in isogenic BTKi^S^ and BTKi^R^ CCMCL1 and Sp49 cells. Erastin (0.5 μM for CCMCL1, 2 μM for Sp49) was used as positive control. (G) GSEA plots for curated Ferroptosis_Suppressors geneset comparing JEKO1^R^ vs. JEKO1^S^ cells (GSE141333) and IB-resistant (patient^R^) vs. –sensitive (patient^S^) MCL patients (GSE141335). All BTKi treatments were performed at 5 μM. Data represent mean ± SD (n=5). Statistical significance was determined by one-way ANOVA with Tukey’s test; ns=non-significant and ***p<0.001, ****p<0.0001.

To examine the contribution of ferroptosis suppression to the development of BTKi resistance in MCL, we next treated isogenic BTKi-sensitive (S) and -resistant (R) CCMCL1 and Sp49 cell lines with BTKi. Remarkably, compared to the robust induction of lipid peroxidation in the sensitive cells, the paired resistant cells exhibited minimal ferroptotic responses (Fig. 1F). Consistently, gene set enrichment analysis (GSEA) of JEKO1^R^ vs. JEKO1^S^ revealed significant enrichment of ferroptosis suppressor gene sets (Fig. 1G, Supplemental Table 1). This enrichment pattern was further validated in primary MCL cells derived from BTKi-resistant vs. -sensitive patients ^41^. Together, these results indicate that ferroptosis suppression is a major mechanism underlying BTKi resistance.

### BRG1 regulates ferroptosis response in MCL cells

As one of the frequently mutated genes in MCL, the SWI/SNF chromatin regulator BRG1 has previously been implicated in clinical resistance to BTKi ^40^. To test the potential contribution of BRG1 mutations to BTKi resistance, we generated paired isogenic cell lines by stably integrating either wild-type or the recurrent T910M mutant BRG1 (Fig. S2A) ^30,42^. Compared with BRG1^WT^-expressing controls, the BRG1^T910M^-expressing MCL cells exhibited notably greater resistance to both covalent (IB, ZB) and reversible (PB) BTKi (Fig. 2A), despite their comparable suppression of BTK (Y233) phosphorylation (Fig. 2B). Importantly, in contrast to BRG1^WT^ cells in which IB treatment led to robust elevation of labile Fe²⁺, ROS, and lipid peroxidation, the paired BRG1^T910M^ cells failed to elicit significant ferroptosis response upon IB treatment (Fig. 2C-E), suggesting resistance to ferroptosis. Introduction of another MCL-associated mutant, BRG1^K785R^ (Fig. S2A), also supported the potential contribution of BRG1 mutations by conferring resistance to IB-induced lipid peroxidation and cell death (Fig. S2B-C). Considering the role of BRG1 as a chromatin remodeler, we hypothesized that aberrant BRG1 may promote BTKi resistance through transcriptionally suppressing ferroptotic responses. Mutant BRG1 alleles thus provide a powerful system to mechanistically dissect the oncogenic consequences of altered BRG1 activity.

**Fig. 2.**
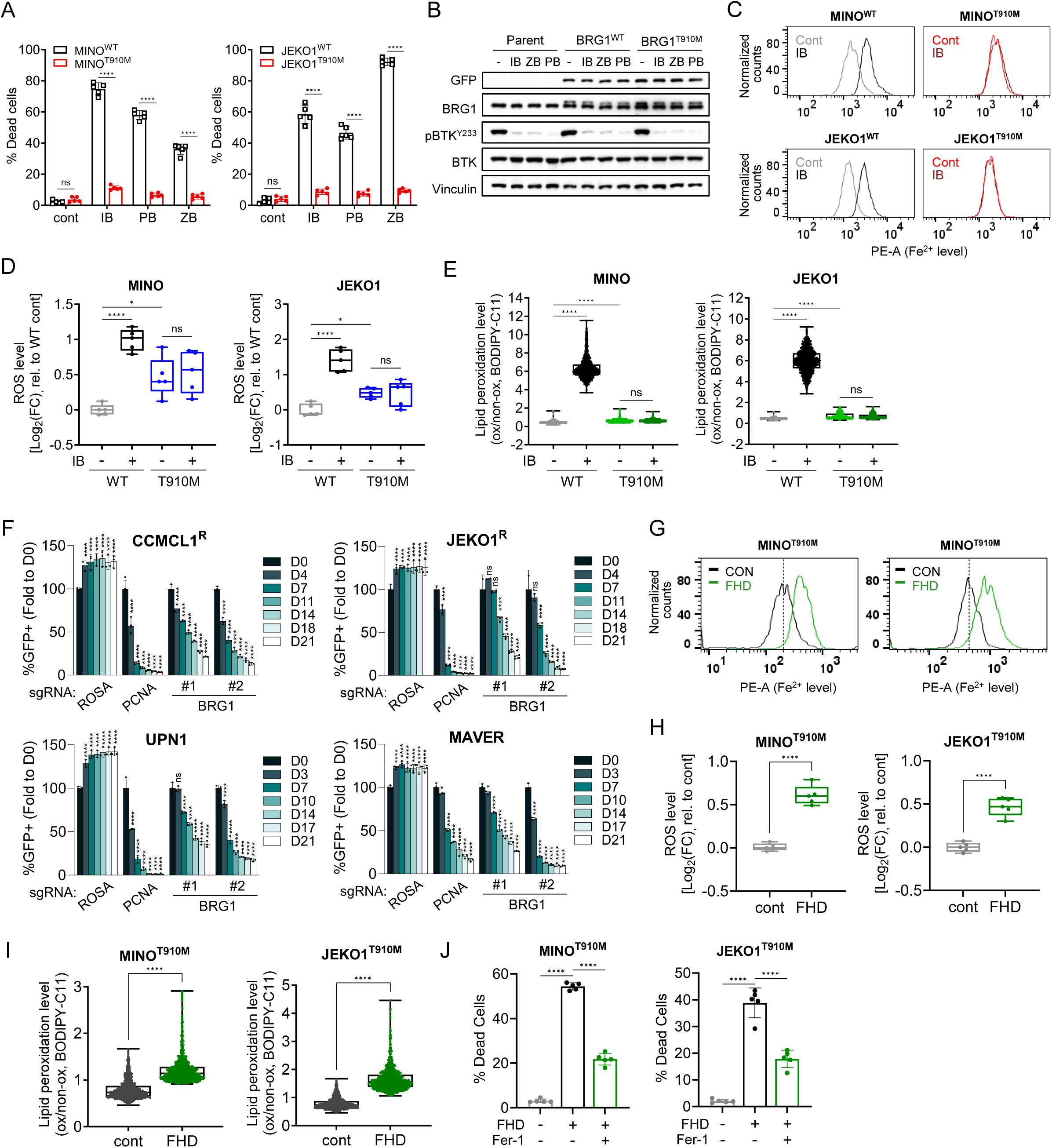
BRG1 regulates ferroptosis response in MCL cells. (A) Cell death response to various BTKi in BRG1^WT^ and BRG1^T910M^ MCL cell lines (96 h). (B) Immunoblot analysis of MINO^Parent^, MINO^WT^, and MINO^T910M^ cells following BTKi treatment (1 h). (C-E) IB-induced changes in labile Fe^2+^ (C; 48 h), ROS (D; 48 h), and lipid peroxidation (E; 72 h) in BRG1^WT^ *vs.* BRG1^T910M^ MCL cells. (F) Competition-based proliferation assays of sgRNAs against BRG1 in Cas9-transduced CCMCL1^R^, MAVER1, JEKO1^R^, and UPN1 cells (n=3). sgROSA was used as a negative control. (G) Representative histograms showing labile Fe^2+^ level changes in BRG1^T910M^ cell lines following FHD-286 treatment (48 h) (H-I) FHD-286-induced changes in ROS (H; 48 h), and lipid peroxidation (I; 72 h) in BRG1^T910M^ MCL cell lines. (J) FHD-286 (96 h)-induced cell death response with or without Fer-1 (2 μM) in BRG1^T910M^ MCL cell lines. All BTKi and FHD-286 treatment were performed at 5 μM and 10 nM, respectively. Unless specified, data represent mean ± SD (n=5). Statistical significance in (H) and (I) was determined using Unpaired t-test; all others used one-way ANOVA with Tukey’s test, ns=non-significant, *p<0.05, **p<0.01, ***p<0.001, ****p<0.0001.

In support of this idea, a CRISPR/Cas9-based vulnerability screen targeting 182 epigenetic regulators with ∼1,461 sgRNAs uncovered that multiple BTKi-resistant MCL cell lines were highly dependent on BRG1 (Fig. S2D, Supplemental Table 2). This dependency was further validated in competition-based proliferation assays using independent BRG1-targeting sgRNAs (Fig. 2F). Moreover, inhibition of BRG1 in BRG1^T910M^-expressing cells by FHD-286, an allosteric BRG1/BRM ATPase inhibitor, induced strong ferroptotic responses, as indicated by the increased labile Fe²⁺ (Fig. 2G), ROS accumulation (Fig. 2H), and elevated lipid peroxidation (Fig. 2I). Importantly, the FHD-286-induced cell death was effectively rescued by co-treatment of ferroptosis inhibitor Fer-1 (Fig. 2J), confirming that its cytotoxicity is mediated through ferroptosis. These findings together indicate that aberrant BRG1 mediates BTKi resistance by transcriptionally suppressing ferroptosis.

### BRG1 regulates ferroptosis through both BTK-dependent and -independent pathways

To investigate the molecular connection between BRG1 and ferroptosis resistance in MCL, we next analyze both nascent and steady-state transcriptional changes by comparing FHD-286- and vehicle-treated JEKO1 and MINO cells. PRO-seq and RNA-seq analyses identified 336 (PRO-seq, |FC|>1.5) and 1,359 DEGs (RNA-seq, |FC|>2.0), respectively (Fig. 3A). Integration of these orthogonal datasets yielded 134 genes consistently regulated upon BRG1 inhibition. KEGG pathway analysis of these core BRG1-regulated genes revealed B-cell receptor signaling (BCR) signaling as a top enriched pathway (enrichment score 5.4; -log_10_ p-value) (Fig. 3B). BRG1 Cut&Run analysis in MINO MCL cells confirmed the BRG1 enrichment at regulatory regions of many genes involved in BCR signaling pathway, including *CD19*, *CD79B*, *RAC2*, *CARD11*, and *SPI1* (Fig. 3C). Further ATAC-seq analysis showed that these BRG1 binding sites largely overlapped with accessible chromatin regions of sgROSA-transduced control cells, which became inaccessible following CRISPR-mediated BRG1 depletion. These findings suggest that BRG1 plays a regulatory role in BCR signaling, likely through direct binding and chromatin remodeling.

**Fig. 3.**
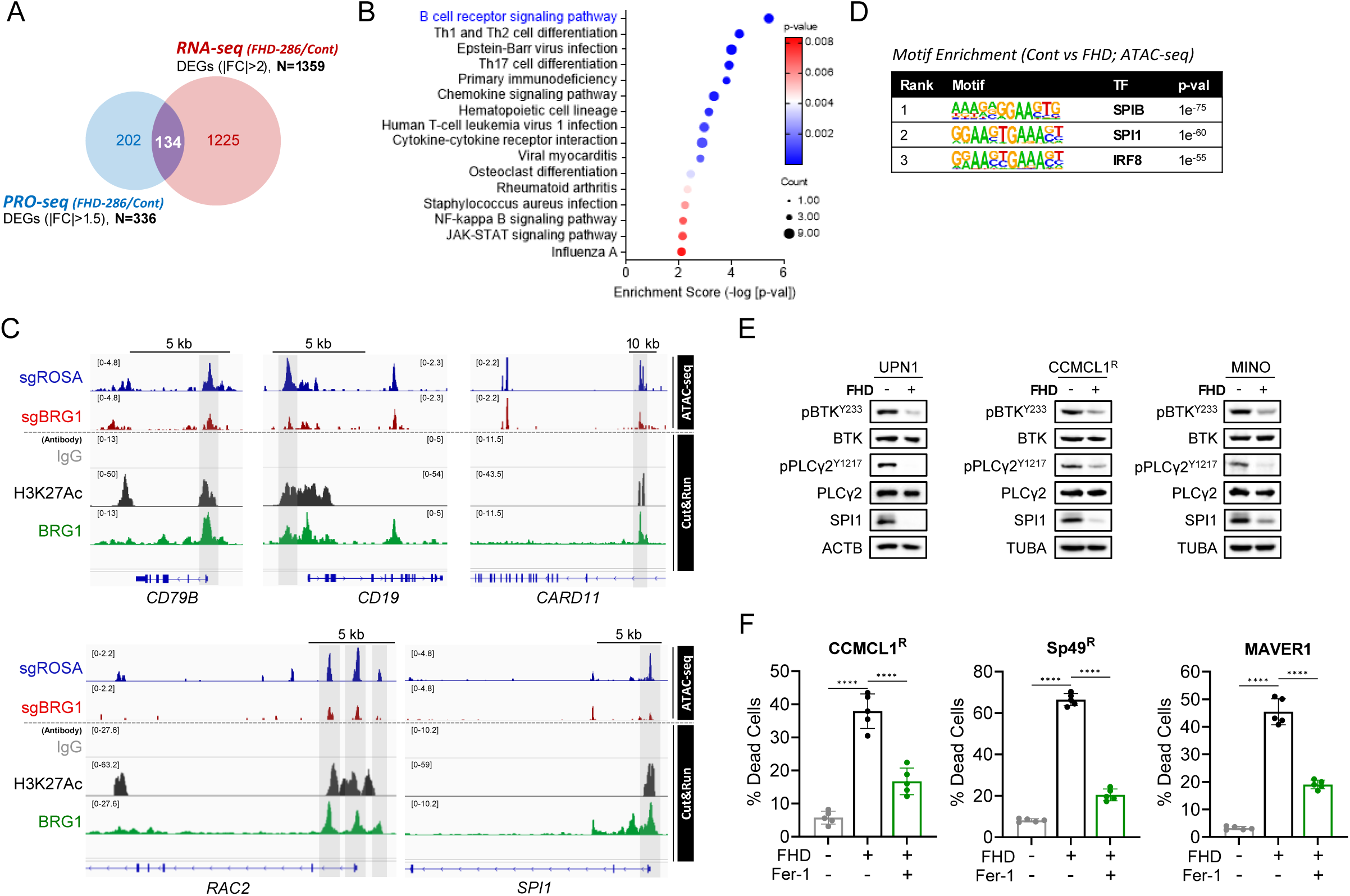
BRG1 regulates ferroptosis through both BTK-dependent and -independent pathways. (A) Venn diagram showing overlap of downregulated genes upon FHD-286 treatment identified by PRO-seq and RNA-seq. (B) KEGG pathway enrichment of the commonly downregulated genes from (A). (C) IGV tracks of ATAC-seq (sgROSA *vs.* sgBRG1 CCMCL1^Δp53^ cells) and IgG, H3K27ac, BRG1 Cut&Run (MINO^WT^ cells) at *CD19*, *CD79B, CARD11, RAC2, and SPI1* loci. (D) HOMER motif enrichment analysis of differentially accessible regions (macs2, q<0.0001) between control and FHD-treated MINO cells (cut-off: p-val<1e^-50^). (E) Immunoblot analysis of multiple MCL cells (UPN1, CCMCL1^R^, MINO) following FHD-286 treatment (10 nM, 48 h). (F) FHD-286 (10 nM, 96 h)-induced cell death response with or without Fer-1 in BTKi resistant MCL cell lines (CCMCL1^R^, Sp49^R^, and MAVER1; n=5, mean ± SD). All statistical significance was determined using one-way ANOVA with Tukey’s multiple-comparison test. ****p<0.0001.

Indeed, HOMER’s motif enrichment analysis of differential ATAC-seq peaks (control *vs.* FHD-286, macs2, q < 0.0001) revealed a significant loss of accessibility upon FHD-286-mediated BRG1 inhibition at the Ets-family motifs, particularly those recognized by the BCR signaling transcription factors *SPI1* (PU.1) and *SPIB* (Spi-B) (Fig. 3D) ^43^. Consistent with this, immunoblot analysis confirmed reduced phosphorylation of BTK and PLCγ2 as well as SPI1 expression following pharmacological BRG1 inhibition in MCL cells (Fig. 3E). Given that BTK signaling protects MCL cells against ferroptosis (Fig. 1, S1B-S1D), these findings suggest that BRG1 regulates ferroptotic responses in MCL through transcriptional control of BCR signaling.

Beyond its role in BCR signaling, our data also indicate that BRG1 may contribute to ferroptosis regulation through additional pathway(s). Specifically, BTKi treatment of resistant MCL cells had minimal effects on ferroptosis induction and cell survival despite effective BTK inhibition (Fig. 1F, 2A-B). By contrast, BRG1 inhibition by FHD-286 triggered significant cell death in all three tested BTKi resistant cell lines (CCMCL1^R^, Sp49^R^, and MAVER1) (Fig. 3F). Notably, this FHD-286-induced cell death was partially rescuable by co-treatment with the ferroptosis inhibitor Fer-1 (Fig. 3F). These results indicate that aberrant BRG1-mediated transcriptional programs can suppress ferroptosis independently of BTK signaling.

### MEF2B mediates ferroptosis response downstream of BRG1

To define the aberrant BRG1-regulated transcriptional programs that promote BTKi resistance in MCL, we conducted RNA-seq comparing isogenic BRG1^T910M^ and BRG1^WT^ cells. The analysis identified 3,127 DEGs (log_2_|FC|>0.6), including 2,237 down- and 890 up-regulated genes in BRG^T910M^ cells (Fig. 4A). To identify direct transcriptional targets, we integrated the RNA-seq data with two ATAC-seq datasets (BRG1^T910M^ *vs* BRG1^WT^; sgROSA *vs.* sgBRG1). The RNA-seq-derived DEG list (log_2_|FC|>1) was first intersected with genes whose chromatin regions exhibited differential accessibility between BRG1^T910M^ and BRG1^WT^ cells (Fig. 4B, blue and orange). The results were subsequently overlapped with genes linked to BRG1-dependent ATAC-seq peaks that lost accessibility upon BRG1 depletion (sgROSA *vs.* sgBRG1) (Fig. 4B, green). This integrative analysis yielded 17 high-confidence BRG1-regulated genes. Among them, *MEF2B* was ranked as the most significantly up-regulated gene in BRG1^T910M^ cells (FDR-adjusted p-value<n.d.) (Fig. 4C). Immunoblot analysis further confirmed that MEF2B protein levels were markedly elevated in BRG1^T910M^ and BRG1^K785R^ cells relative to paired wild-type controls (Fig. S3A, S3B).

**Fig. 4.**
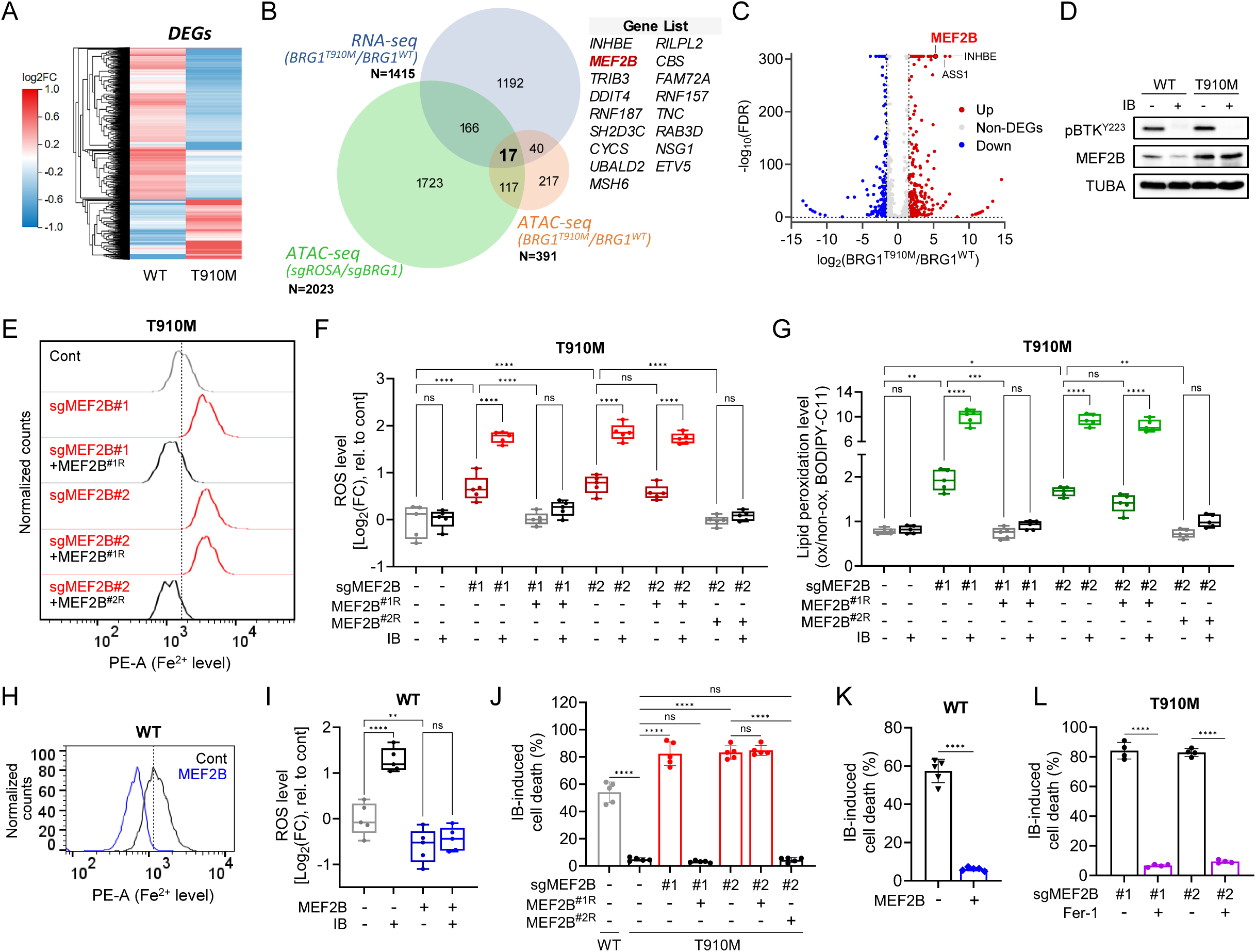
MEF2B mediates ferroptosis response downstream of BRG1. (A) Heatmap of DEGs from RNA-seq comparing BRG1^T910M^ and BRG1^WT^ MINO cells. Rows represent individual genes and columns represent samples, with fold changes shown relative to parental MINO controls. (B) Venn diagram showing overlap between DEGs (RNA-seq: MINO^T910M^ *vs.* ^WT^) and differentially accessible regions (ATAC-seq: CCMCL1^Δp53^ sgROSA *vs.* sgBRG1). (C) Volcano plot of RNA-seq data comparing MINO^T910M^ and ^WT^ cells, which each point representing an individual gene. (D) Immunoblot of JEKO1^WT^ and JEKO1^T910M^ cells following IB treatment (5 μM, 24 h) (E-G) MEF2B depleted and rescued JEKO1^T910M^ cells showing basal labile Fe^2+^ (E), IB-induced ROS (F; 48 h), and lipid peroxidation (G; 72 h) level changes. (H-I) MEF2B-overexpressed JEKO1^WT^ cells showing basal Fe^2+^ (H) and IB-induced ROS (I; 48 h) level changes. (J-K) IB-induced cell death (96 h) in MEF2B-depleted and rescued JEKO1^T910M^ cells (J) and MEF2B-overexpressed JEKO1^WT^ cells (K). (L) Effect of Fer-1 (2 μM) on IB-induced cell death in MEF2B-depleted BRG1^T910M^ cells (96 h). All IB treatments were performed at 5 μM. Data represent mean ± SD (n=5). Statistical significance was determined by one-way ANOVA with Tukey’s test; ns=non-significant, *p<0.05, **p<0.01, ***p<0.001, ****p<0.0001.

Notably, IB treatment markedly reduced MEF2B protein expression in wild-type but not mutant BRG1-expressing MINO and JEKO1 cells (Fig. 4D, S4A), supporting the idea that aberrant BRG1-mediated transcriptional activity promotes BTKi resistance by suppressing ferroptosis. Consistently, CRISPR-mediated MEF2B depletion significantly elevated basal labile Fe²⁺ levels in BRG1^T910M^ MINO and JEKO1 MCL cells (Fig. 4E, S4B, and S4C). Moreover, MEF2B depletion in BRG1^T910M^ cells also elevated both baseline and BTKi-induced ROS accumulation and lipid peroxidation, all of which were fully suppressed by CRISPR-resistant MEF2B re-expression, ruling out off-target effects (Fig. 4F, 4G, S4B, and S4D-S4F). Conversely, ectopic MEF2B expression in BRG1^WT^ cells reduced basal labile Fe^2+^ levels (Fig. 4H, S4G, and S4H) as well as both baseline and BTKi-induced ROS levels (Fig. 4I, S4G, S4I, and S4J), indicating that MEF2B indeed mediates BRG1-associated ferroptotic response.

Given our findings of MEF2B as a downstream effector of the BRG1-mediated transcriptional axis, we next investigated its role in BTKi resistance. As expected, MEF2B depletion sensitized BRG1^T910M^-expressing MCL cells to BTKi-induced cell death, an effect that was fully reversible upon CRISPR-resistant MEF2B re-expression (Fig. 4J, S4B, S4K, and S4L). Importantly, this sensitization could also be abrogated by co-treatment with the ferroptosis inhibitor Fer-1 (Fig. 4K, S4M, and S4N). Conversely, ectopic MEF2B expression in BRG1^WT^ cells conferred resistance to BTKi-induced cell death, further supporting its key role in BRG1-associated ferroptotic suppression (Fig. 4L, S4O, and S4P).

### NDUFA4L2 serves as the key downstream effector of MEF2B to mediate BTKi-induced ferroptosis resistance

MEF2B is a transcription factor crucial for normal B-cell development ^44^. To elucidate the mechanism by which BRG1-MEF2B axis promotes ferroptosis resistance, we first identified 2,378 DEGs (log_2_|FC|>0.6) from RNA-seq analysis of BRG1^WT^ cells ectopically expressing MEF2B (Fig. 5A, green). These were enriched for direct MEF2B targets by intersecting with 1,857 genes associated with MEF2B Cut&Run peaks (Fig. 5A, red). We further refined this set by intersecting with 1,415 DEGs from BRG1^T910M^ *vs.* BRG1^WT^ RNA-seq (log_2_|FC|>1), identifying 32 overlapping candidates (Fig. 5A, blue, gene list). Among these, we focused on *NDUFA4L2* and *FTH1* based on their reported roles in ferroptosis-relevant processes: redox metabolism ^45^ and iron storage ^46^, respectively (Fig. 5A). Cut&Run revealed MEF2B occupancy at acetylated histone H3 lysine 27 (H3K27Ac)-positive regulatory regions of both *FTH1* and *NDUFA4L2* loci (Fig. 5B). Consistently, BRG^T910M^-expressing cells showed markedly elevated protein levels of MEF2B, NDUFA4L2, and FTH1 in (Fig. 5C). While MEF2B depletion reduced expression of both NDUFA4L2 and FTH1 (Fig. 5D, S5A), NDUFA4L2 depletion did not affect MEF2B levels (Fig. S5B), establishing MEF2B as an upstream regulator of both genes.

**Fig. 5.**
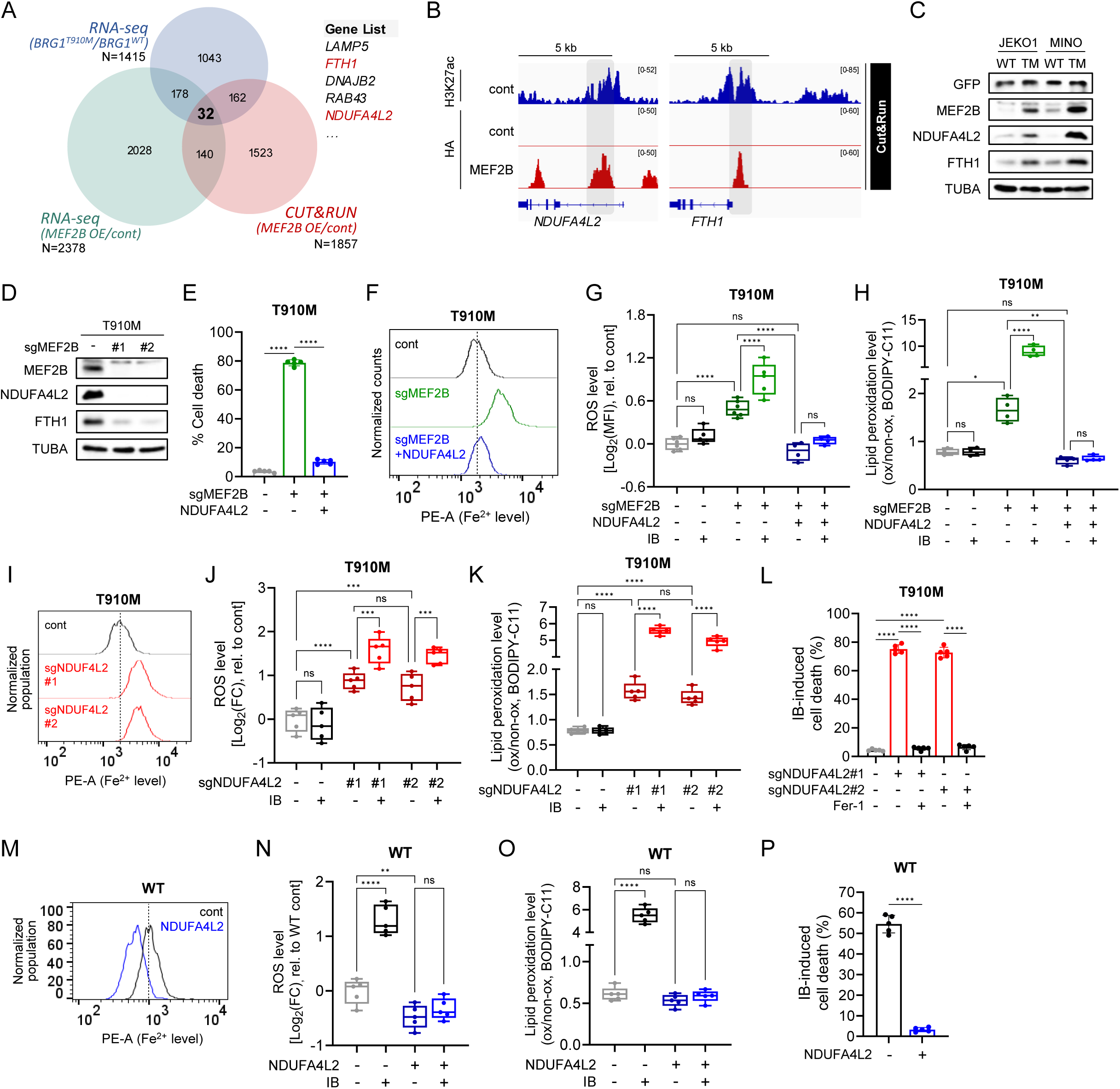
NDUFA4L2 serves as the key downstream effector of MEF2B to mediate BTKi-induced ferroptosis resistance. (A) Venn diagram showing overlap of DEGs from RNA-seq (MINO BRG1^T910M^ *vs.* ^WT^; JEKO1 MEF2B OE *vs.* control), and HA-MEF2B Cut&Run. (B) IGV tracks of HA-MEF2B and H3K27ac Cut&Run at the *NDUFA4L2* and *FTH1* loci. (C-D) Immunoblot analysis of BRG1^WT^ *vs.* BRG1^T910M^ MCL cells (C) and MEF2B-depleted JEKO1^T910M^ MCL cells (D). (E) IB-induced cell death (96 h) in MEF2B-depleted JEKO1^T910M^ cells with or without NDUFA4L2 rescue. (F-H) MEF2B-depleted JEKO1^T910M^ cells with or without NDUFA4L2-reconstitution showing basal labile Fe^2+^ (F), IB-induced ROS (G; 48 h), and lipid peroxidation (H; 72 h) level changes. (I-L) NDUFA4L2-depleted JEKO1^T910M^ cells showing basal Fe^2+^ (I), IB-induced ROS (J; 48 h), and lipid peroxidation (K; 72 h) level changes. (L) Effect of Fer-1 (2 μM) on IB-induced cell death in NDUFA4L2-depleted JEKO1^T910M^ cells (96 h). (M-O) NDUFA4L2-overexpressed JEKO1^WT^ cells showing basal Fe^2+^ (M), IB-induced ROS (N; 48 h), and lipid peroxidation (O; 72 h) level changes. (P) IB-induced cell death in NDUFA4L2-overexpressed JEKO1^WT^ cells (96 h). All IB treatments were performed at 5 μM. Data represent mean ± SD (n=5). All statistical significance was determined by one-way ANOVA with Tukey’s test; ns=non-significant, *p<0.05, **p<0.01, ***p<0.001, ****p<0.0001.

To assess the role of NDUFA4L2 in BRG1/MEF2B-driven ferroptosis suppression, we performed rescue experiments by ectopically expressing NDUFA4L2 in MEF2B-depleted BRG1^T910M^ cells (Fig. S5C). NDUFA4L2 reconstitution restored resistance to IB, counteracting the sensitizing effects induced by MEF2B loss (Fig. 5E, S5D). It also reversed the phenotypic consequences of MEF2B depletion, restoring both baseline and IB-induced increases in labile Fe²⁺, ROS and lipid peroxidation to levels comparable with those of control BRG1^T910M^ cells (Fig. 5F-5H, S5E-S5G). These results identify NDUFA4L2 as a key downstream effector of MEF2B that mediates BRG1-driven resistance to BTKi-induced ferroptosis.

Consistently, NDUFA4L2 depletion in BRG1^T910M^ MCL cells markedly elevated basal labile Fe²⁺ (Fig. 5I, S5H), as well as both baseline and IB-induced ROS (Fig. 5J, S5I), and lipid peroxidation (Fig. 5K, S5J). This depletion also sensitized the cells to IB-induced ferroptosis (up to 75% for JEKO1; 68% for MINO), which was largely reversed by co-treatment with the ferroptosis inhibitor Fer-1 (down to ∼6% for both JEKO1 and MINO) (Fig. 5L, S5K). Conversely, ectopic overexpression of NDUFA4L2 in BRG1^WT^ MCL cells suppressed basal free Fe²⁺ (Fig. 5M, S5L), as well as both baseline and BTKi-induced ROS (Fig. 5N, S5M) and lipid peroxidation (Fig. 5O, S5N), thereby reducing IB-induced cell death (Fig. 5P, S5O). Altogether, these findings establish NDUFA4L2 as a key downstream effector of the BRG1-MEF2B axis that mediates ferroptosis resistance.

### NDUFA4L2 regulates mitochondrial activity to promote ferroptosis resistance

As a mitochondrial protein, NDUFA4L2 mitigates oxidative stress by inhibiting Complex I ^45^. In BRG1^WT^ cells, ectopic expression of NDUFA4L2 suppressed mitochondrial biogenesis, turnover, and oxidative phosphorylation, as evidenced by reduced mitochondrial mass (Fig. 6A, S6A), decreased mitophagy (Fig. 6B, S6B), and a ∼4.6-fold reduction in mitochondrial oxygen consumption rate (Fig. 6C, 6D). Conversely, NDUFA4L2 depletion in BRG1^T910M^ cells significantly increased mitochondrial mass (Fig. 6E, S6A), mitophagy (Fig. 6F, S6B), and oxygen consumption (∼1.7-fold increase; Fig. 6G, 6H), suggestive of enhanced oxidative phosphorylation and more active clearance of damaged mitochondria.

**Fig. 6.**
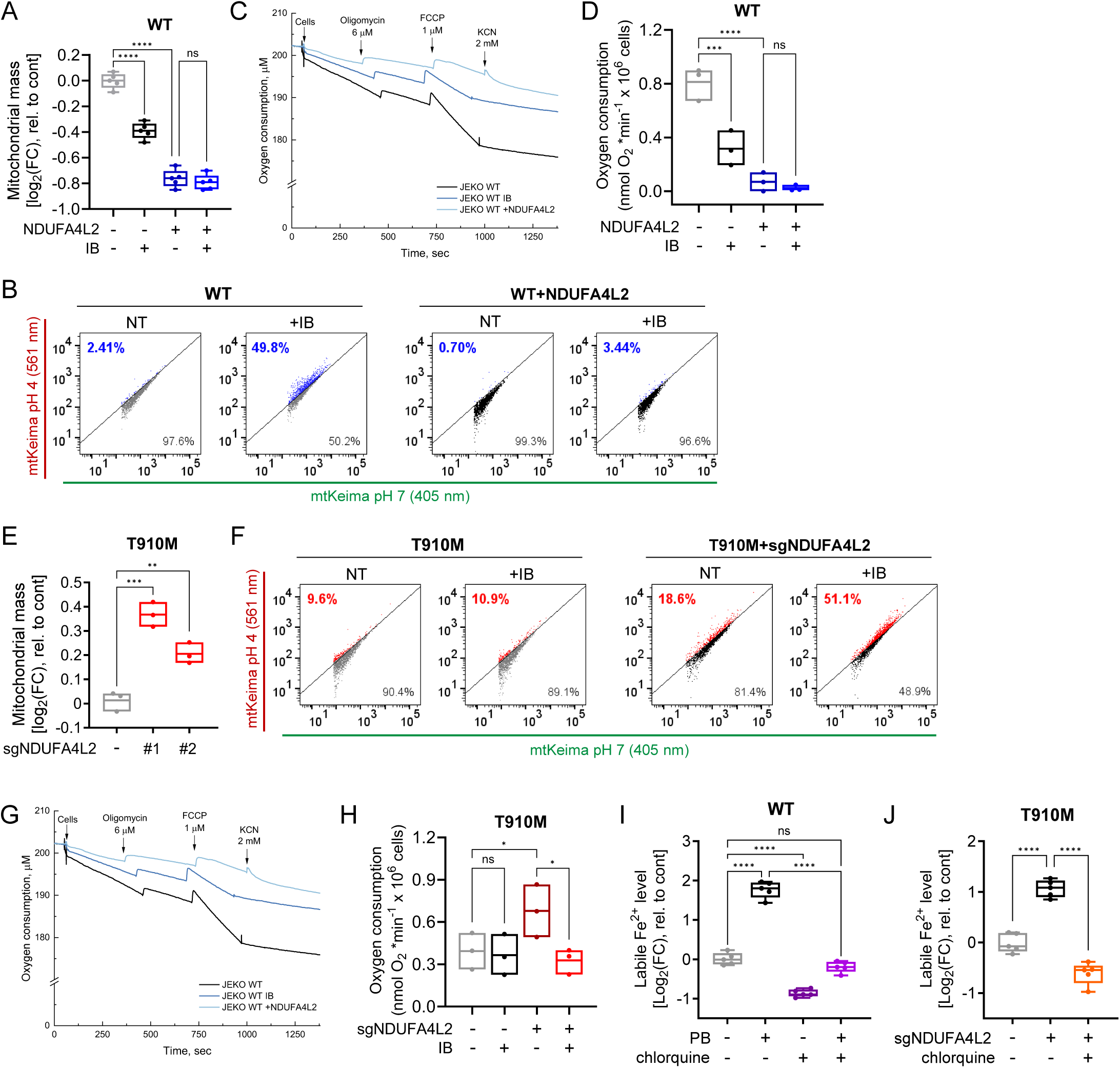
NDUFA4L2 regulates mitochondrial activity to promote ferroptosis resistance. (A-D) Relative mitochondrial mass (A; n=5), mitophagy level (B; representative scatter plots), oxygen consumption trace (C), and total OCR (D; n=3) in control *vs.* NDUFA4L2-overexpressed JEKO1^WT^ cells with or without IB treatment (24 h). (E-H) Relative mitochondrial mass (E; n=3), mitophagy level (F; representative scatter plots), oxygen consumption trace (G) and total OCR (H; n=3) in control *vs.* NDUFA4L2-depleted JEKO1^T910M^ cells with or without IB treatment (24 h). (I-J) Labile Fe^2+^ levels in: (I) JEKO^WT^ cells treated with PB and/or chloroquine (100 μM), and (J) NDUFA4L2-depleted JEKO^T910M^ cells treated with chloroquine (100 μM) (24 h; n=5). All IB/PB treatment was performed at 5 μM. Data represent mean ± SD. Statistical significance in (D) and (H) was determined using one-way ANOVA with Uncorrected Fisher’s LSD multiple-comparison test; all others used one-way ANOVA with Tukey’s test, ns=non-significant, *p<0.05, **p<0.01, ***p<0.001, ****p<0.0001.

We further observed that BTK depletion reduced mitochondrial mass (Fig. S6C), consistent with previous reports that BTK inhibition induces mitochondrial dysfunction ^47,48^. In line with this, IB treatment in BRG1^WT^ cells elevated ROS levels (Fig. 4I, 5N) and mitophagy (Fig. 6B, S6B), while reducing mitochondrial respiration (∼2.6-fold; Fig. 6C, 6D) and mitochondrial mass (Fig. 6A, S6A). Notably, these changes were accompanied by suppressed endogenous NDUFA4L2 protein expression (Fig. S6D). Ectopic expression of NDUFA4L2 largely blocked these IB-induced changes (Fig. 5N, 6A-6D, S6A, S6B), supporting its protective role against mitochondrial dysfunction. Conversely, depletion of NDUFA4L2 in BRG1^T910M^ MCL cells, which are BTKi-resistant and express elevated endogenous NDUFA4L2, further enhanced mitophagy upon IB treatment (Fig. 6F, S6B), and sensitized these cells to IB-induced reduction in oxygen consumption (Fig. 6G, 6H), resembling the phenotype observed in BRG1^WT^ cells. These findings indicate that NDUFA4L2 protects MCL cells from IB-induced mitochondrial damage by shifting metabolism away from oxidative phosphorylation.

Ferroptosis depends on the balance of redox status and labile iron availability, both of which are closely linked to mitochondrial function ^49^. Notably, PB-induced elevation of labile Fe²⁺ was blocked by co-treatment with the autophagy inhibitor chloroquine (Fig. 6I, S6E). Similarly, NDUFA4L2 depletion–induced Fe²⁺ accumulation in BRG1^T910M^ cells was fully suppressed by chloroquine (Fig. 6J, S6F), indicating that mitochondrial damage–associated mitophagy is the primary source of free Fe²⁺ in these settings. Together, these findings support our hypothesis that NDUFA4L2, as a downstream effector of the BRG1–MEF2B axis, protects MCL cells from BTKi-induced ferroptosis by limiting mitochondrial oxidative metabolism and restricting mitophagy-associated ROS and iron release.

### BRG1 inhibition re-sensitizes resistant MCL cells to BTKi

The BRG1 inhibitor FHD-286 is currently in a Phase I trial for relapsed or refractory acute myeloid leukemia and myelodysplastic syndrome ^50,51^. Given our findings that BRG1 is a crucial regulator of ferroptosis in MCL, we next evaluated the therapeutic potential of pharmacological BRG1 inhibition against this disease. Indeed, FHD-286 as a single agent significantly suppressed the growth of all nine MCL cell lines tested, independent of their BTKi resistance status, with IC₅₀ values uniformly in the nanomolar range (Fig. 7A). Moreover, FHD-286 exhibited robust cytotoxic effects in six primary cultures derived from independent IB-resistant MCL patients, highlighting its potential to overcome BTKi resistance (Fig. 7B). Notably, MCL cell lines harboring BRG1^T910M^ as well as the patient sample with M781I mutation of BRG1 both exhibited greater sensitivity to FHD-286 compared to those with wild-type BRG1 (Fig. 7A, 7B, S7A, S7B). These findings suggest that BRG1 mutations may enhance cellular dependency on BRG1 activity, potentially through alternative resistance mechanisms independent of BTK.

**Fig. 7.**
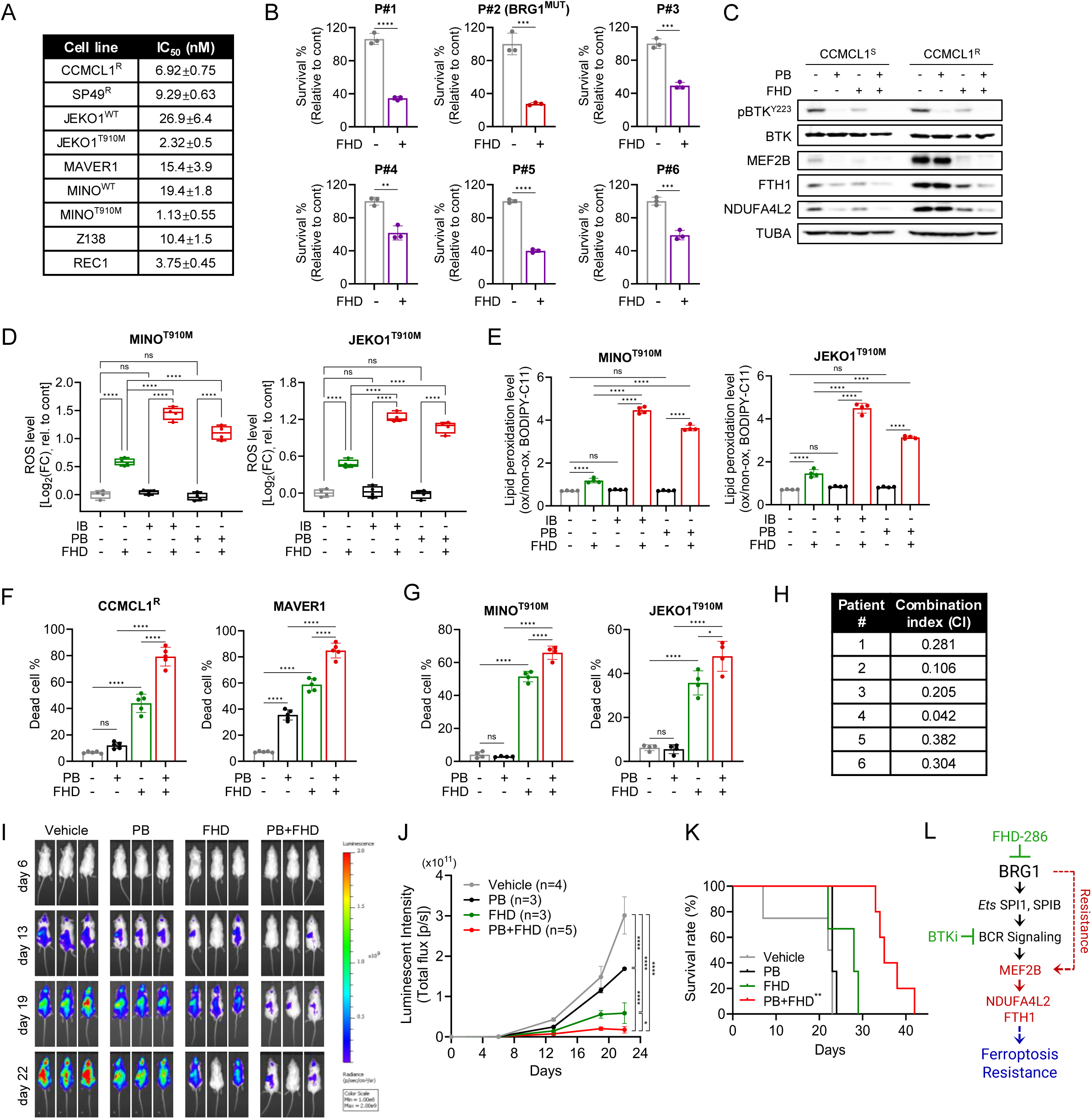
BRG1 inhibition re-sensitizes resistant MCL cells to BTKi. (A) IC50 value of FHD-286 against different MCL cell lines (96 h, n=3). (B) Cytotoxic effect of FHD-286 on primary MCL cultures (96 h, n=3). (C) Immunoblot of CCMCL1^S^ and ^R^ cells treated with PB, FHD-286, or their combinations for 24 h. (D-E) Changes in ROS (D; 48 h, n=5) and lipid peroxidation (E; 72 h n=5) following treatment with IB, PB, FHD-286 or their combinations. (F-G) Cell death effect of FHD-286 (10 nM) and PB (5 μM) combination in BTKi^R^ (CCMCL1^R^, MAVER1; F) and BRG1^T910M^ (MINO^T910M^, JEKO1^T910M^; G) MCL cells (96 h, n=5). (H) CI value for IB and FHD-286 in primary MCL cultures. (I) Representative bioluminescent images from CCMCL1^R^ CDX mice treated with PB (10 mg/kg; n=3), FHD-286 (1.2 mg/kg; n=3), or their combination (n=6). Mean ± SEM. (J) Quantification of bioluminescent images from (I). (K) Kaplan-Meier survival curves of mice from (I-J). Statistical significance was determined using log-rank test (**p=0.0014). (L) Schematic summary of demonstrated pathways in the study. All BTKi and FHD-286 treatment for in vitro and ex vivo experiments were performed at 5 μM and 10 nM, respectively. Unless specified, data represent mean ± SD, and statistical significance was determined using one-way or two-way ANOVA with Tukey’s multiple-comparison test. ns=non-significant, ***p<0.001, ****p<0.0001.

FHD-286 as a single agent induced cell death in BTKi-resistant MCL cells, but was insufficient to fully suppress their growth, achieving only partial therapeutic efficacy (Fig. 2J, 3F). While BRG1 regulates both BTK-dependent and -independent pathways, treatment of CCMCL1^S^ and CCMCL1^R^ cells with FHD-286 only partially suppressed BTK activity (Fig. 7C), which may explain its limited efficacy. This prompted us to evaluate whether combining FHD-286 with BTKi could achieve more complete BTK pathway suppression and enhanced therapeutic efficacy. Remarkably, combined treatment of FHD-286 with pirtobrutinib (PB), a reversible BTKi with improved target coverage and pharmacokinetics ^52^, not only fully inhibited the BTK signaling, it also enhanced suppression of the MEF2B-NDUFA4L2/FTH1 signaling axis in BTKi-resistant CCMCL1 as well as MINO^T910M^ and JEKO1^T910M^ cells (Fig. 7C, Fig. S7C). Consequently, the combination treatment induced robust ferroptosis, shown by elevated ROS production (Fig. 7D), lipid peroxidation (Fig. 7E), and markedly enhanced cell death (Fig. 7F, 7G, S7D). Similarly, a competition-based proliferation assay revealed that BTKi treatment accelerated the selective elimination of BRG1-depleted cells (Fig. S7E). The synergistic cytotoxicity of the combined treatment was further confirmed in primary MCL cultures derived from the IB-resistant patients, with combination index values ranging from 0.042 to 0.382 (Fig. 7H, S6F), suggesting that BRG1 inhibition re-sensitizes BTKi-resistant MCL cells to BTK inhibition.

We next assessed the therapeutic potential of this combination in vivo using a CCMCL1^R^ cell-derived xenograft (CDX) model (Fig. 7I). As expected, bioluminescence imaging revealed that while FHD-286 monotherapy achieved only moderate tumor growth suppression, PB alone elicited minimal effect compared with vehicle controls. By contrast, combined FHD-286/PB treatment produced pronounced anti-tumor activity (Fig. 7J). Consistent with these tumor burden results, survival curves revealed little to no benefit with PB monotherapy and a modest improvement with FHD-286 (median survival extended from 22.5 to 28 days). In comparison, the FHD-286/PB combination markedly prolonged survival, extending the median to 35 days (p=0.0040, Fig. 7K). Collectively, these findings indicate that combining FHD-286 and BTKi effectively suppresses the MEF2B-NDUFA4L2/FTH1 axis and restores ferroptotic sensitivity in BTKi-resistant MCLs (Fig. 7L). These data provide a preclinical rationale for combined BTKi and BRG1 inhibition as a therapeutic strategy in refractory MCL.

## DISCUSSION

Dynamic transcriptional and metabolic rewiring is a key driver of therapy resistance. Here, we identify aberrant BRG1-driven ferroptosis resistance mediated through MEF2B-NDUFA4L2/FTH1 expression. As a key mediator of resistance, MEF2B directly induces the transcription of NDUFA4L2 and FTH1, suppressing mitochondrial metabolism and iron levels against BTKi-induced ferroptosis. We further uncovered that BRG1 inhibition suppresses MEF2B-NDUFA4L2/FTH1 gene expression network thereby promoting ferroptosis. Accordingly, combined BRG1 and BTK inhibition synergistically suppresses MCL progression, supporting BRG1 targeting as a strategy to overcome BTKi-resistance. These findings reveal a key role for BRG1 in regulating ferroptosis susceptibility, establishing a mechanistic link between epigenetic dysregulation and BTKi resistance.

In MCL, aggressive subsets show elevated iron metabolism ^53,54^ posing them more vulnerable to ferroptotic stress. While BTKi are known to induce primarily apoptosis, our findings identify ferroptosis as a complementary mechanism of BTKi-induced cytotoxicity, particularly in TP53-mutant MCL, where apoptotic signaling is compromised. Induction of ferroptosis was consistently observed across multiple BTKi with distinct modes of target engagement (Fig. 1B-1D) and was reproduced by BTK depletion (Fig. S1B-S1D), indicating an on-target effect. Notably, ferroptosis-associated changes were abrogated in MCL cells under aberrant BRG1-driven transcription program (Fig. 2C-2E, S2C), but were restored by FHD-286 treatment (Fig. 2G-2J). These findings show that BRG1 mediates BTKi resistance by suppressing ferroptosis which is a key vulnerability of MCL.

MEF2B is a transcription factor implicated in GC-derived lymphomas, where it regulates BCL6 and is frequently mutated to enhance oncogenic activity ^44,55^. In MCL, MEF2B mutations are less common (∼3-9%) ^56,57^, and its oncogenic role remains largely uncharacterized. Our study identifies MEF2B as a previously unrecognized effector of ferroptosis resistance, upregulated by aberrant BRG1 activity (Fig. 4, S3-S4). Unlike the MYC-driven oxidative phosphorylation program, where MEF2B cooperates with MYC and DNMT3A to promote IB resistance by enhancing oxidative phosphorylation ^58^, BRG1 mutations induce MEF2B expression in a distinct regulatory context that bypasses MYC and reprograms resistance through ferroptosis suppression. These findings highlight MEF2B as a converging node in BTKi resistance, enabling engagement of mechanistically distinct survival pathways.

NDUFA4L2 reduces oxidative phosphorylation and oxygen consumption by inhibiting mitochondrial Complex I activity ^45,59^. It is up-regulated for metabolic adaptation by HIF1α under hypoxia and oxidative stress, where it facilitates metabolic adaptation by limiting ROS production. We identified NDUFA4L2 as a direct downstream effector of MEF2B which serves as an upstream regulator of mitochondrial respiration (Fig. 5), protecting cells against BTKi-caused mitochondrial damage (Fig. 6A-6H, S6A, S6B). This metabolic shift restricts the pool of dysfunctional mitochondria reducing mitophagy-mediated lysosomal degradation (Fig. 6B, 6F, S6B) and release of Fe^2+^ from iron sulfur clusters (Fig. 6I, 6J, S6E, S6F). Together with MEF2B-induced FTH1 (Fig. 5B, 5D, S5A), NDUFA4L2 limits iron availability for lipid peroxidation, collectively suppressing BTKi-induced ferroptosis. Thus, through identification of the BRG1-MEF2B-NDUFA4L2/FTH1 axis, our results link epigenetic dysregulation to metabolic reprogramming as a driver of drug resistance.

While mutant BRG1 promotes BTKi resistance through activation of the MEF2B-NDUFA4L2/ FTH1 axis, we also identify BRG1 as an essential gene, as its complete loss compromises MCL cell viability. Mechanistically, BRG1 sustains pro-survival BCR signaling by maintaining widespread chromatin accessibility at Ets transcription factor binding sites, particularly those of SPI1 and SPIB (Fig. 3D). This dual role of BRG1 partially aligns with the findings from Deng et al. ^36^, which described BRG1 as a haploinsufficient tumor suppressor in GC-derived lymphomas, fine-tuning centrocyte fate through regulating SPI1, IRF family members, and NFκB. In their model, complete BRG1 loss impaired GC formation, whereas heterozygosity facilitated lymphomagenesis.

The mechanism of BRG1 mutation driven resistance to IB and venetoclax combination was demonstrated by BRG1 knockdown which reduced the chromatin accessibility at the *ATF3* locus and led to increased expression of the anti-apoptotic protein BCL-xL ^40^. Notably, our mutant BRG1 ectopic expression model uncovered a distinct mechanism, characterized by the upregulation of MEF2B-NDUFA4L2 (Fig. 4-5. S3-S5). These divergent mechanisms may act synergistically to enhance resistance, as both ultimately converge on inhibiting cell death. Together, these findings underscore the need for precision medicine approaches, as different BRG1 alterations (loss vs. mutation) can drive resistance through distinct yet complementary pathways.

Consistent with the prior report that ARID1A-mutant lymphomas exhibit enhanced sensitivity to BAF complex inhibition ^60^, our study demonstrates that BRG1 mutant MCL cells are more susceptible to FHD-286, supporting a synthetic lethal interaction in BTKi-refractory disease. Mechanistically, FHD-286 inhibits both SPI1/SPIB-driven BCR signaling and MEF2B expression, collectively promoting ferroptosis (Fig. 7L). These findings provide a strong mechanistic rationale for dual targeting of BRG1 and BTK, which resulted in significant synergistic cytotoxicity across multiple preclinical MCL models (Fig. 7F–7K, S7D, S7F). Altogether, targeting BRG1-dependent resistance to ferroptosis represents a promising therapeutic strategy for BTKi-refractory MCL.

## Methods

Information about the antibodies, chemical and biologicals, cell lines, oligonucleotides, recombinant DNA, software, algorithms, and instruments are included under Supplemental Table 3.

### Cell lines and culture details

MCL cell lines CCMCL1, JEKO1, UPN1, MAVER1, MINO, Z138, and REC1 were cultured in RPMI1640 supplemented with 10% heat-inactivated fetal bovine serum (FBS), 2 mM L-glutamine, and 50 unit/ml penicillin-streptomycin in a 37 ℃ humidified 5% CO2 incubator. All cell lines used for gene knockdown were lentivirally transduced to express human codon optimized SpCas9 nuclease for establishment of somatic deletion cells by CRISPR/Cas9 genome editing. Infected cells were selected with blasticidin or puromycin for 7-10 days. BTKi-resistant CCMCL1^R^ and Sp49^R^ cell line were generated as previously described ^41^, by maintaining them in gradually increasing concentration of BTKi (i.e., 10 μM IB, ZB, or PB) over 8 weeks. The BRG1^WT^ and BRG1^T910M^-expressing paired isogenic cell lines were generated using the PiggyBac transposon system delivered by electroporation for stable genomic integration, as previously described ^61^. Both pPB CAG SMARCA4-Venus (Addgene plasmid # 153950) and pPB CAG SMARCA4 T910M-Venus (Addgene plasmid # 153951) were gifts from Luca Tiberi (http://n2t.net/addgene:153950 and http://n2t.net/addgene:153951; RRID:Addgene_153950 and RRID:Addgene_153951). All cell lines were repeatedly tested for mycoplasma contamination by PCR.

### Primary MCL ex vivo treatment

Peripheral blood from IB-resistant, relapse MCL patients was obtained following written informed consent under a protocol approved by the Institutional Review Board of The Ohio State University in accordance with the Declaration of Helsinki. MCL cells were isolated using CD19 magnetic beads and purity was determined by flow cytometry analysis using CD45, CD5, and CD20 staining. Primary MCL cells with TP53 mutations were cultured in RPMI 1640 medium supplemented with 20% FBS, 50 U/mL penicillin-streptomycin, Glutamax, IL-6 (40 U/mL), soluble IL-6R (40 U/mL), IL-10 (50 ng/mL), IGF-1 (30 ng/mL), sCD40L (0.5 μg/mL), and BAFF (50 ng/mL). A round bottom 96 well plate with 150 μL of 2x10^6^/mL cells were treated with drugs for 72-96 h and the number of live cells was determined.

### sgRNA and expression constructs

All the sgRNAs were cloned into BsmB1-digested LRG2.1 or lenticrisprV2 vectors. pXPR_016-sgATF4-1 was a gift from William Hahn (Addgene plasmid # 202455 ; http://n2t.net/addgene:202455 ; RRID:Addgene_202455). For the ectopic expression of HA-MEF2B, a full length MEF2B with N-terminal HA tag was amplified with using CloneAmp™ HiFi PCR Premix and cloned into pLU vector by Gibson assembly.

### Cytotoxic killing analysis and viability assay

For cytotoxic killing analysis, 5.0x10^4^ MCL cells in 2 mL were treated with compounds. After 4 days, the fraction of dead cells was determined by FACS analysis for % of ToPro3-staining positive cells. For IC50 of FHD-286, cell viability was assessed using the XTT assay. Briefly, 1 × 10⁴ cells/well were seeded in 96-well plates and treated with the indicated doses for triplicate. After 96 hours, XTT labeling reagent mixed with electron coupling reagent was added directly to each well and incubated 3 h at 37°C. Formation of formazan dye was quantified by absorbance at 450 nm with a reference wavelength of 650 nm using a microplate reader.

### Measuring cellular ROS, labile iron, or lipid peroxidation levels

CellROX™ Deep Red Reagent (Thermo Fisher Scientific, C10422) was used at 0.5 μM for 1 h incubation in growth media to detect oxidative stress and ROS levels. Oxidized probe fluorescence was measured by flow cytometry at excitation/emission (ex/em) maxima of ∼644/665 nm. Labile iron levels were assessed using the FerroOrange fluorescence probe (Cell Signaling Technology, 36104S). Cells were harvested, washed once with HBSS and incubated with 1 μM probe for 30 min at 37 °C, followed by measurement of fluorescence intensity at ex/em maxima of ∼540/580 nm. Intracellular lipid peroxidation was evaluated using BODIPY-C11 (Cayman Chemical, #27086). Cells were incubated at 5 μM probe in growth medium for 1 h, and fluorescence was detected by flow cytometry at ex/em maxima of 581/591 nm (non-oxidized) and 488/510 nm (oxidized). The ratio of oxidized to non-oxidized signal was calculated to quantify lipid peroxidation levels.

### Mitophagy and mitochondria mass measurement

Mitophagy was measured using mt-Keima ratiometric probe. pHAGE-mt-mKeima was a gift from Richard Youle (Addgene plasmid # 131626 ; http://n2t.net/addgene:131626; RRID:Addgene_131626). Cells were lentivirally infected to express mt-mKeima. Mitophagy levels were determined by acquiring intensity of emission spectra using BV605 (violet laser 405 nm) together with PE-CF594 (yellow laser 565 nm) and emission spectra with 610/20 BP. MitoTracker® Deep Red FM (Thermo Fisher Scientific) probe was used to measure mitochondrial mass. Cells were harvested, washed with PBS, and incubated with 50 nM MitoTracker ® Deep Red FM in pre-warmed serum-free medium at 37°C for 30 minutes. Flow cytometry was performed to determine the mean fluorescence intensity (MFI) which was used as a proxy for mitochondrial mass.

### ATAC-seq analysis

ATAC-seq was performed at the Cornell University BRC Epigenomics Facility (RRID:SCR_021287). Briefly, 55,000 cells flash-frozen in 10% DMSO were lysed, permeabilized, and tagmented using the Omni ATAC-seq protocol ^62^ followed by 12 cycles of barcoding PCR. Gel-purified libraries were sequenced on the Element Biosciences AVITI to obtain approximately 10M paired-end reads.

### Cleavage under targets and release using nuclease (Cut&Run) analysis

50,000 MCL cells were harvested and fixed with 0.1% formaldehyde for 5 min and further processed by Epicypher ChIC system using validated antibodies. All the recovered DNA was used for dual indexed library preparation and sequenced for 10M reads per sample at Weill Cornell Genomics Core.

### ATAC-seq and Cut&Run data analysis

Sequencing reads were aligned to hg38 reference genome using Bowtie v2.4.4 with the parameter (-sensitive --no-unal -p 20 -x). Peak calling was performed by MACS2 v1.4.2 with default parameters. BigWig files and analysis of read density in peak regions were generated using deeptools v3.5.1. Read density of specific genomic regions were displayed using Integrative Genomics Viewer (IGV) v2.17.2. To identify enriched transcription factor binding motifs, we performed motif discovery using HOMER (v4.11) on differentially accessible chromatin regions. Differential peaks identified by comparing MACS2-called narrowPeak files were used as input for HOMER’s findMotifsGenome.pl script. Motif enrichment was calculated based on known vertebrate motif databases provided with HOMER, and results were ranked by statistical significance (hypergeometric p-value).

### RNA-seq analysis

Total RNAs were isolated from cells using NucleoSpin RNA kit and subjected to RNA sequencing at the Genomics Resources Core facility of Weill Cornell Medicine. RNA-seq libraries were prepared using the Illumina TruSeq stranded mRNA library preparation kit and sequenced on HiSeq4000 sequencer (Illumina). Before analysis, library-quality was assessed using fastQC and adapters/sequencing artifacts were removed using cutadapt (v1.8.2). Gene expression quantification was accessed with selective-alignment and fragment/GC content bias correction against the GENCODE 41 human transcripts annotations, using salmon (v1.6.0). The expression counts were normalized by the trimmed mean of M-values (TMM) method and transformed to counts per million.

### Gene Set Enrichment analysis (GSEA)

GSEA was performed using the GSEA software (GSEA v4.4.0, Broad Institute) ^63^ on publicly available datasets GSE141331 and GSE141335, with samples grouped by IB responsiveness. For GSE141335, only patients showing concordance between clinical response and ex vivo IB testing were included (9 resistant, 13 sensitive). A ferroptosis suppressor gene set was curated from FerrDb V2 ^64^, restricted to genes with TPM ≥5 across MCL cell lines (Supplemental Table 1). Analyses were run with 1,000 permutations, and gene sets with nominal p<0.05 and FDR q<0.1 were considered significantly enriched.

### PRO-seq analysis

Chromatin or cells were incubated in the nuclear run-on reaction condition (5 mM Tris-HCl pH 8.0, 2.5 mM MgCl2, 0.5 mM DTT, 150 mM KCl, 0.5% Sarkosyl, 0.4 units/μL of RNase inhibitor) with biotin-NTPs and rNTPs supplied (18.75 μM rATP, 18.75 μM rGTP, 1.875 μM biotin-11-CTP, 1.875 μM biotin-11-UTP for uPRO; 18.75 μM rATP, 18.75 μM rGTP, 18.75 μM rUTP, 0.75 μM CTP, 7.5 μM biotin-11-CTP for pChRO) for 5 min at 37°C. Run-On RNA was extracted using TRIzol, and fragmented under 0.2 N NaOH for 15 min on ice. Fragmented RNA was neutralized, and buffer exchanged by passing through P-6 columns (Biorad). 3′ RNA adaptor (/5Phos/NNNNNNNNAGAUCGGAAGAGCACACGUCUG/ 3InvdT/) is ligated at 5 μM concentration for 1 hours at room temperature using T4 RNA ligase (NEB), followed by 2 consecutive streptavidin bead bindings and extractions. Extracted RNA is converted to cDNA using template switch reverse transcription with 1 μM RT primer (GTGACTGGAGTTCAGACGTGTGCTCTTCCGATC), 3.75 μM Template Switch Oligo (TCTTTCCCTACACGACGCTCTTCCGATCTrGrGrG), 1x Template Switch Enzyme and Buffer (NEB) at 42°C for 30 min. After a SPRI bead clean-up, the cDNA is PCR amplified up to 20 cycles using primers compatible with Illumina TRU-seq sequencing.

### KEGG pathway enrichment analysis

KEGG pathway enrichment analysis was performed using the DAVID Bioinformatics Resources ^65^ online platform. A total of 134 overlapping DEGs from PRO-seq (|FC| > 1) of JEKO1 cells and RNA-seq (|FC| > 1.5) of MINO cells, each comparing FHD-286-treated vs. control, were analyzed. Enriched pathways were identified using default DAVID parameters, with those showing p < 0.01 considered significant and presented in the plot.

### Pirtobrutinib and FHD-286 treatment of MCL xenografts

BTKi-resistant CCMCL1^R^ cells were lentivirally transduced to express firefly luciferase. Two million cells were tail vein injected into NSG (NOD.Cg-PrkdcscidIl2rgtm1Wjl/SzJ) mice that were randomized into four groups: vehicle (1% DMSO in 20% 2-hydroxypropyl-β-cyclodextrin), pirtobrutinib, 10 mg/kg, QD), FHD-286 (1.2 mg/kg, QD), or combination therapy. Treatments were done by i.p. injection for a 3-5 week period. Tumor growth was monitored via weekly bioluminescent imaging with an IVIS Spectrum system (Caliper Life Sciences). Use of laboratory mice were approved by the Weill Cornell Medicine Institutional Animal Care and Usage Committee (IACUC protocol 2018-0039)

### Calculation of combination index (CI)

Combination index (CI) values were calculated using CompuSyn software (ComboSyn Inc.) based on the Chou–Talalay method, using cell viability percentages from ex vivo MCL primary cell cultures treated with FHD-286, IB, or their combination. CI < 1, = 1, and > 1 indicated synergism, additivity, and antagonism, respectively.

### Statistical analysis

We determined experimental sample sizes based on preliminary data. For numerical variables, all results are expressed as mean ± STDEV. Normal distribution of the sample sets was determined before applying two-sample Student’s t-test or one-way ANOVA for group comparisons. Unless specified, a two-sample Student’s t-test was used to compare the differences between two groups and the one-way ANOVA was used to assess the differences among multiple groups followed by a post-hoc with Tukey’s procedure to adjust the p-values for multiple comparisons. Kaplan-Meier method was used to estimate the probability of survival, and the log-rank test was used to compare the overall survival difference between groups. All statistical tests are two-sided, and the differences were considered significant when p<0.05. GraphPad Prism (ver. 10) software was used for all statistical analyses.

## Data availability

The data that support the findings of this study are available within the article and its Supplemental Tables. RNA-seq, PRO-seq, ATAC-seq and Cut&Run data can be found on Gene Expression Omnibus (GEO) database with accession number GSE305144, GSE303985.

## Acknowledgements

The authors are deeply grateful to the MCL patients who generously provided tissue samples for this study, and to the Ohio State University Comprehensive Cancer Center Leukemia Tissue Bank Shared Resource (supported by NCI P30-CA016058) for assistance with sample procurement. We also thank Foghorn Therapeutics for providing FHD-286. This work was supported in part by a P01 grant (CA214274) and an R01 grant (CA276349) from the National Institutes of Health/National Cancer Institute, by a Mantle Cell Lymphoma Research Initiative grant from the Leukemia and Lymphoma Society (MCL7001–18), and by the National Research Foundation of Korea (RS-2024-00411768). SYH was additionally supported by the Basic Science Research Program through the National Research Foundation of Korea, funded by the Korean government (Ministry of Science and ICT)(RS-2024-00414906).

## Contributions

S.-Y.H., H.Z., and J.P. conceived and designed the study. S.-Y.H., H.N., H.K., and J.P. developed methodology. S.-Y.H., H.N., B.Y.-S, S.Y., I.H., H.K., and J.P. acquired data. S.-Y.H., B.Y.-S, A.G., H.K., and J.P. analyzed and interpreted data (e.g., statistical analysis, computational analysis). S.- Y.H., H.Z., and J.P. wrote the manuscript. L.S., L.A., R.A.B., X.H., M.DL., S.C-K., H.Z., and J.P., provided administrative, technical, or material support, and reviewed the data. H.Z., and J.P. supervised the study.

---

**Fig. S1.** Ferroptosis suppression underlies BTK inhibitor resistance. (A) Effect of Fer-1 (2 μM) on PB or ZB-induced (96 h) cell death in MCL cells. (B-D) Changes in labile Fe^2+^ (B), ROS (C; n=3) and lipid peroxidation (D; n=5) following BTK depletion in MCL cells (6-day post-sgRNA treatment). All BTKi treatments were performed at 5 μM. Data represent mean ± SD. Statistical significance was determined by one-way ANOVA with Tukey’s test; ***p<0.001, ****p<0.0001.

**Fig. S2.** BRG1 regulates ferroptosis response in MCL cells. (A) Schematic diagrams summarizing the distribution of mutations along the BRG1 sequence (http://oncokb.org). Mutation diagram circles are colored with respect to the corresponding mutation types. (B) Cell death response to various BTKi in BRG1^WT^ and BRG1^K785R^ MCL cell lines (96 h). (C) IB-induced changes in lipid peroxidation (72 h) in BRG1^WT^ *vs.* BRG1^K785R^ MCL cells. (D) Ranked CRISPR/Cas9-based viability screening in CCMCL1^R^, JEKO1^R^, and UPN1. All IB treatments were performed at 5 μM. Data represent mean ± SD. Statistical significance was determined by one-way ANOVA with Tukey’s test; ns=non-significant, ****p<0.0001.

**Fig. S3.** Elevated MEF2B in BRG1^mut^-expressing MCL cells. (A-B) Immunoblot analysis of BRG1^WT^ *vs.* BRG1^T910M^ (A) or BRG1^K785R^ (B) MCL cells.

**Fig. S4.** MEF2B mediates ferroptosis response downstream of BRG1. (A) Immunoblot of MINO^WT^ and MINO^T910M^ cells following IB treatment (24 h). (B) Immunoblot of MEF2B-depleted and rescued JEKO1^T910M^ and MINO^T910M^ cells. (C-D) MEF2B-depleted and rescued MINO^T910M^ cells showing basal labile Fe^2+^ (C; 48 h) and IB-induced ROS (D; 48 h) level changes. (E) PB-induced ROS (48 h) level changes in MEF2B-depleted/rescued JEKO1^T910M^ and MINO^T910M^ cells. (F) IB-induced lipid peroxidation (72 h) level changes in MEF2B-depleted/rescued MINO^T910M^ cells. (G) Immunoblot of MEF2B-overexpressed JEKO1^WT^ and MINO^WT^ cells. (H-I) MEF2B-overexpressed MINO^WT^ cells showing basal labile Fe^2+^ (H; 48 h) and IB-induced ROS (I; 48 h) level changes. (J) PB-induced ROS (48 h) level changes in MEF2B-overexpressed JEKO1^WT^ and MINO^WT^ cells. (K) IB-induced cell death (96 h) in MEF2B-depleted/rescued MINO^T910M^ cells (L) PB-induced cell death (96 h) in MEF2B-depleted/rescued JEKO1^T910M^ and MINO^T910M^ cells. All IB/PB treatments were performed at 5 μM. Data represent mean ± SD (n=5). Statistical significance was determined by one-way ANOVA with Tukey’s test; ns=non-significant, *p<0.05, **p<0.01, ***p<0.001, ****p<0.0001.

**Fig. S5.** NDUFA4L2 serves as the key downstream effector of MEF2B to mediate BTKi-induced ferroptosis resistance. (A-C) Immunoblot of MEF2B-depleted MINO^T910M^ cells (A), NDUFA4L2-depleted MINO^T910M^ and JEKO1^T910M^ cells (B), and MEF2B-depleted MINO^T910M^ and JEKO1^T910M^ cells reconstituted with NDUFA4L2 (C). (D) IB-induced cell death (96 h) in MEF2B-depleted MINO^T910M^ cells with or without NDUFA4L2 rescue. (E-G) MEF2B-depleted MINO^T910M^ cells with or without NDUFA4L2-reconstitution showing basal labile Fe^2+^ (E), IB-induced ROS (F; 48 h), and lipid peroxidation (G; 72 h) level changes. (H-J) NDUFA4L2-depleted MINO^T910M^ cells showing basal Fe^2+^ (H), IB-induced ROS (I; 48 h), and lipid peroxidation (J; 72 h) level changes. (K) Effect of Fer-1 (2 μM) on IB-induced cell death in NDUFA4L2-depleted MINO^T910M^ cells (96 h). (L-N) NDUFA4L2-overexpressed MINO^WT^ cells showing basal Fe^2+^ (L), IB-induced ROS (M; 48 h), and lipid peroxidation (N; 72 h) level changes. (O) IB-induced cell death in NDUFA4L2-overexpressed MINO^WT^ cells (96 h). All IB treatments were performed at 5 μM. Data represent mean ± SD (n=5). Statistical significance was determined by one-way ANOVA with Tukey’s test; ns=non-significant, *p<0.05, ***p<0.001, ****p<0.0001.

**Fig. S6.** NDUFA4L2 regulates mitochondrial activity to promote ferroptosis resistance. (A-B) Relative mitochondrial mass (A) and mitophagy levels (B) in MINO^WT^ (control *vs.* NDUFA4L2-overexpressed; n=5) and MINO^T910M^ (control vs. NDUFA4L2-depleted; n=3) cells with or without IB treatment (24 h). (C) Relative mitochondrial mass in BTK-depleted MCL cells (6-day post sgRNA transduction). (D) Immunoblot of MINO^WT^ and MINO^T910M^ cells with or without IB treatment (24 h). (E-F) Labile Fe^2+^ levels in: (E) MINO^WT^ cells treated with PB and/or chloroquine (100 μM), and (F) NDUFA4L2-depleted MINO^T910M^ cells treated with chloroquine (100 μM) (24 h; n=5). All IB/PB treatment was performed at 5 μM). Data represent mean ± SD. Statistical significance with two experimental group was determined using Unpaired t-test; Besides, one-way ANOVA with Tukey’s test was applied; ns=non-significant, *p<0.05, **p<0.01, ***p<0.001, ****p<0.0001.

**Fig. S7.** BRG1 inhibition re-sensitizes resistant MCL cells to BTKi. (A) Dose-response curve of FHD-286 in multiple MCL cell lines (96 h, n=3). (B) Dose-response curve comparing FHD-286-induced cell death in BRG1^WT^ vs. BRG1^T910M^ or ^K785^ MCL cell lines (96 h, n=5). (C) Immunoblot of BRG1^WT^ and BRG1^T910M^ cells treated with either IB, FHD-286, or their combinations for 24 h. (D) Cell death effect of FHD-286 (10 nM) and IB (5 μM) combination in BRG1^T910M^ MCL cell lines. (E) Competition-based proliferation assays of sgBRG1 in Cas9-transduced CCMCL1^R^, MAVER1 cells with or without PB (2 μM) (n=3). sgROSA was used as a negative control. Two-way ANOVA with Holm-Sidak’s multiple-comparison. (F) Relative cell survival rate following treatment with IB (5 μM), FHD-286 (10 nM), or their combinations in primary MCL cultures (n=3). Data represent mean ± SD. Unless specified, significance was determined using one-way ANOVA with Tukey’s multiple-comparison test. ns=non-significant, ***p<0.001, ****p<0.0001.

