## Supplementary figures and images for "Leveraging BRG1 Driven Ferroptosis Resistance to Overcome Treatment Resistance"

### Supplemental Figures

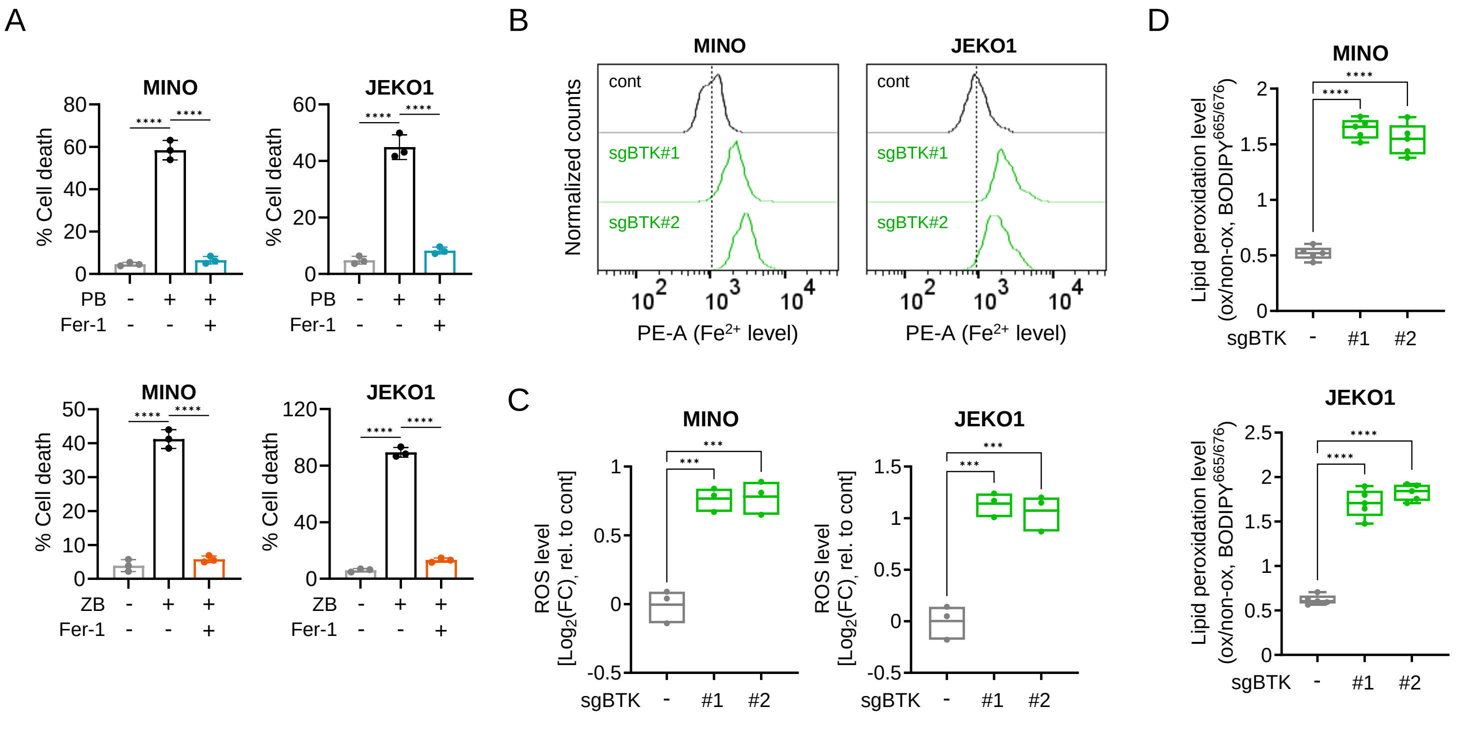

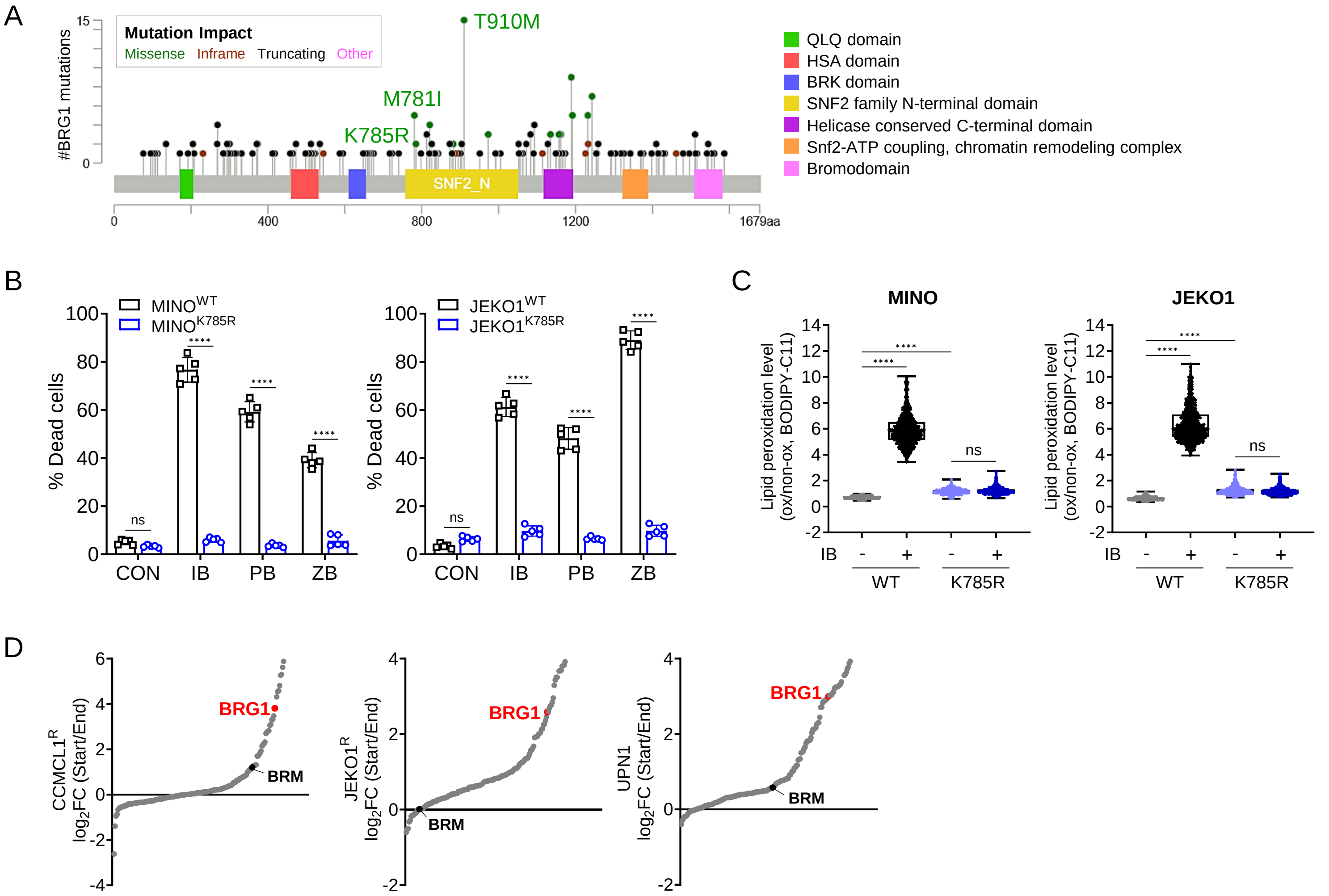

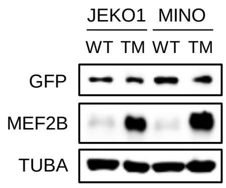

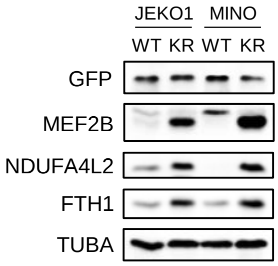

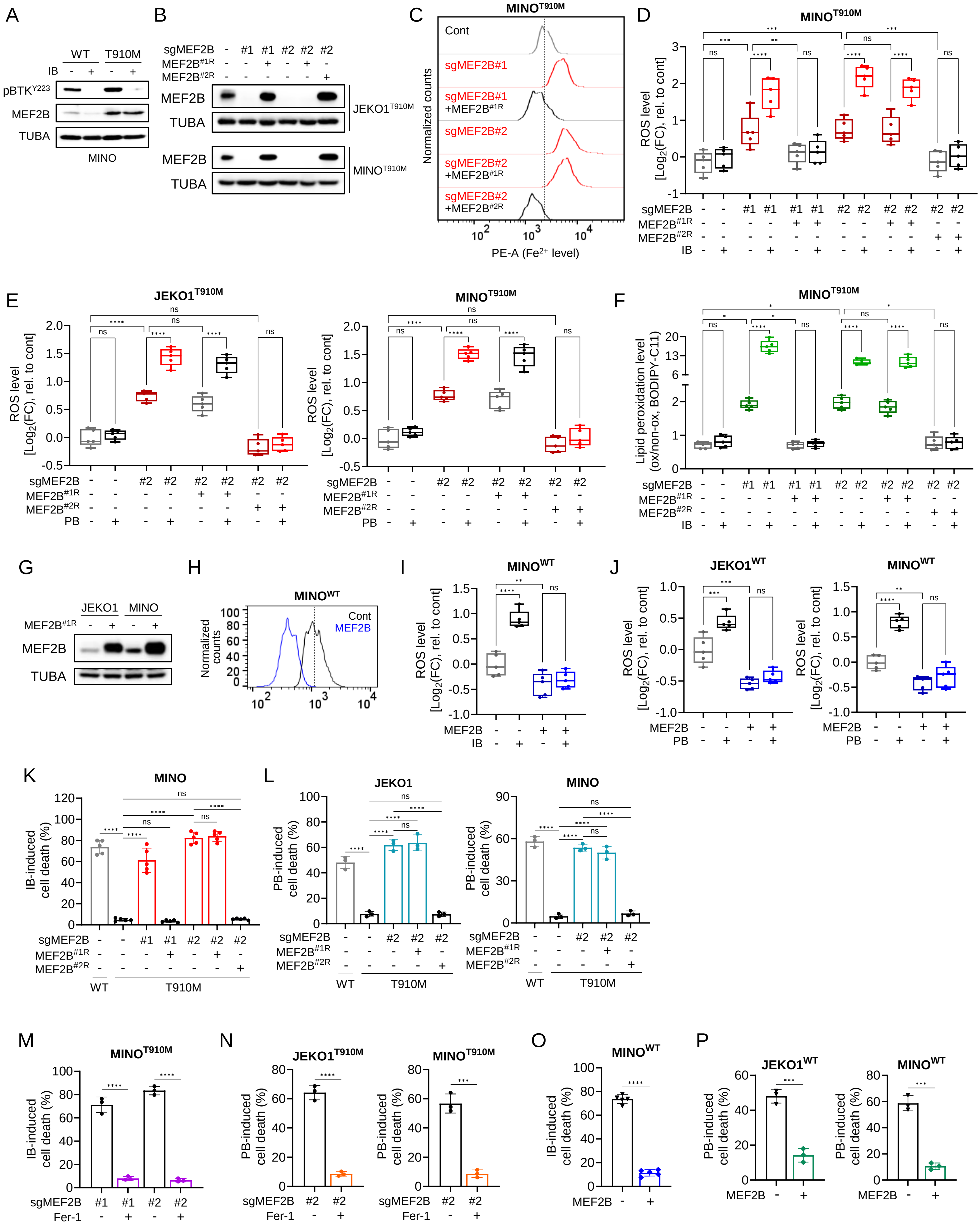

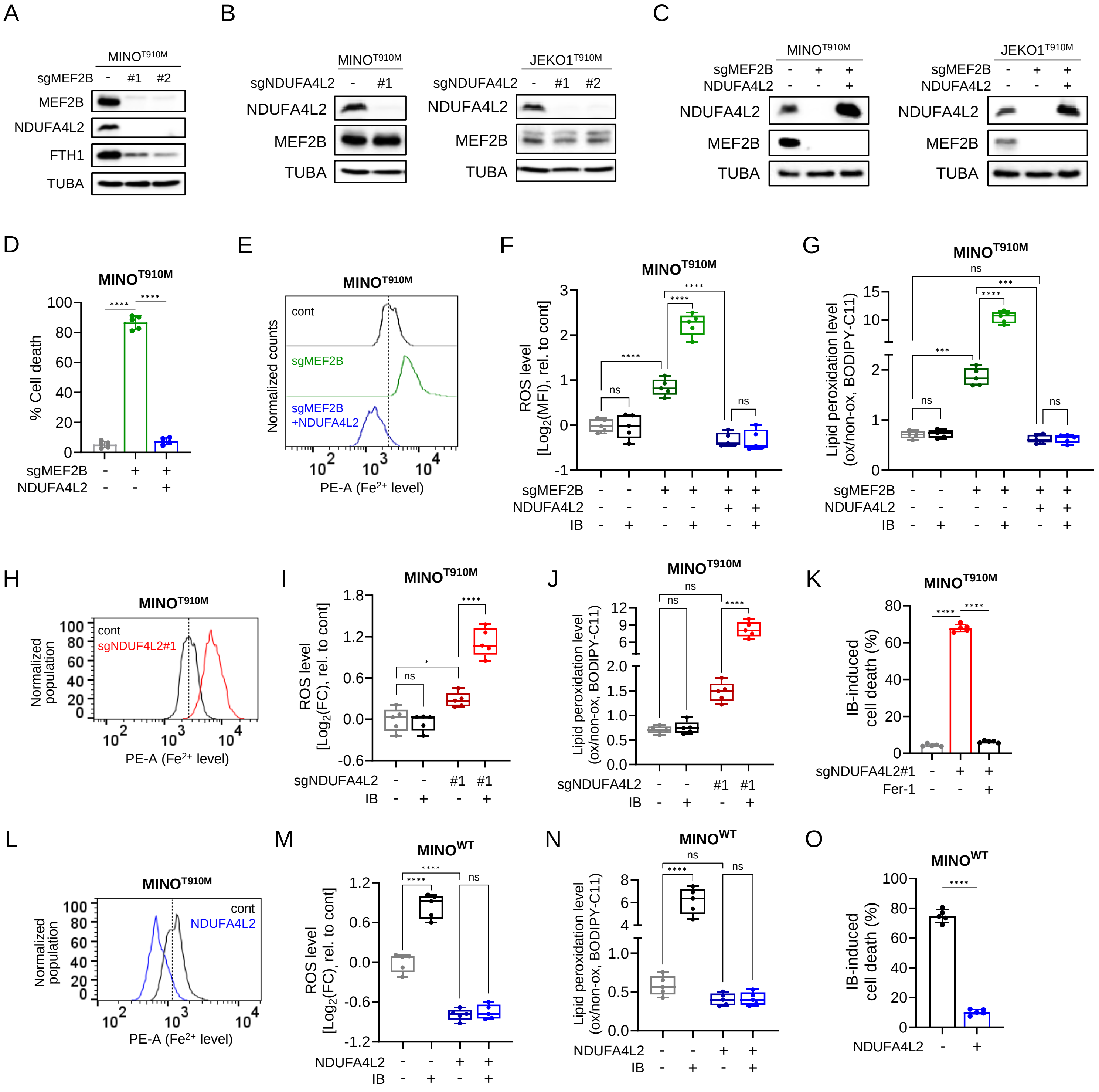

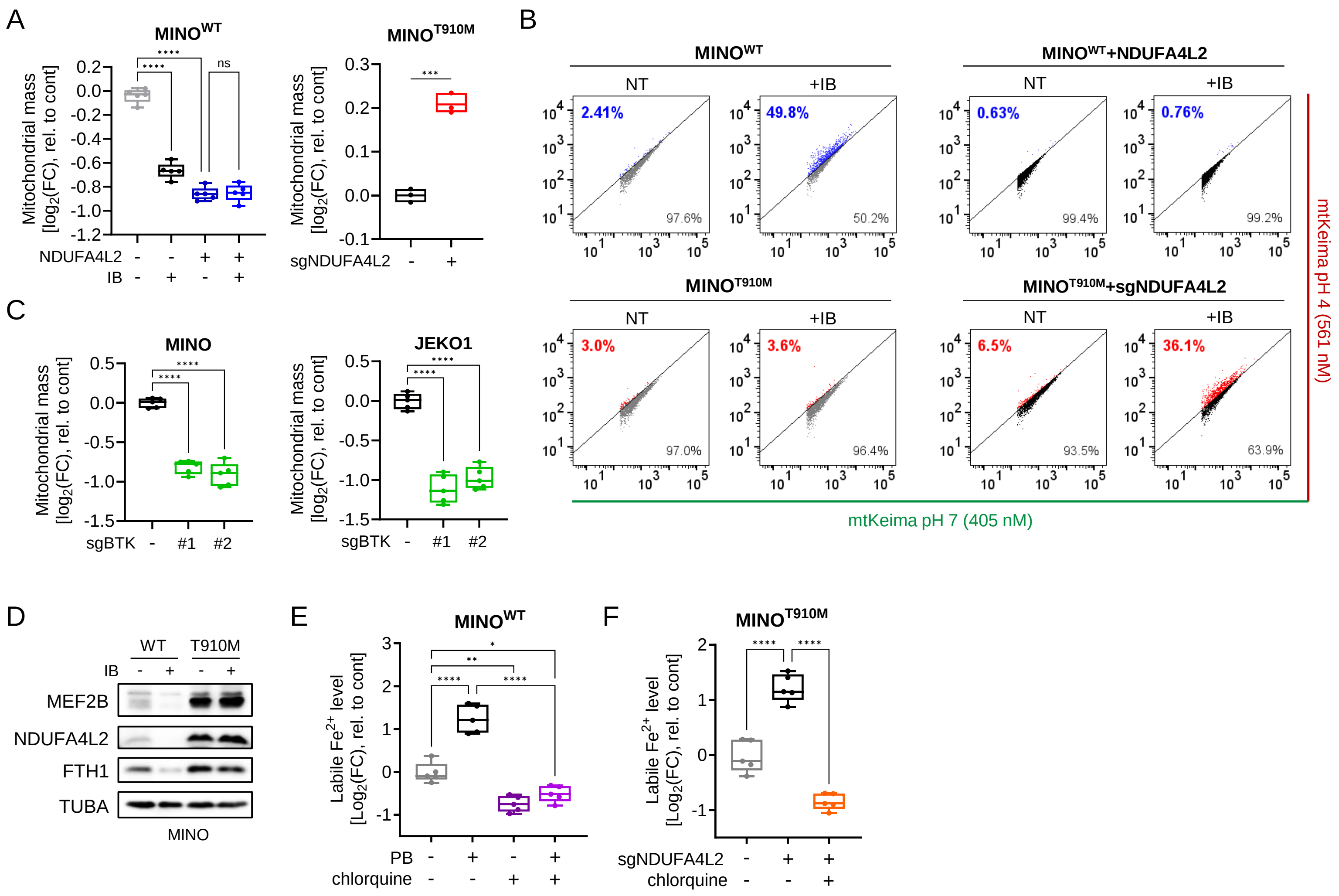

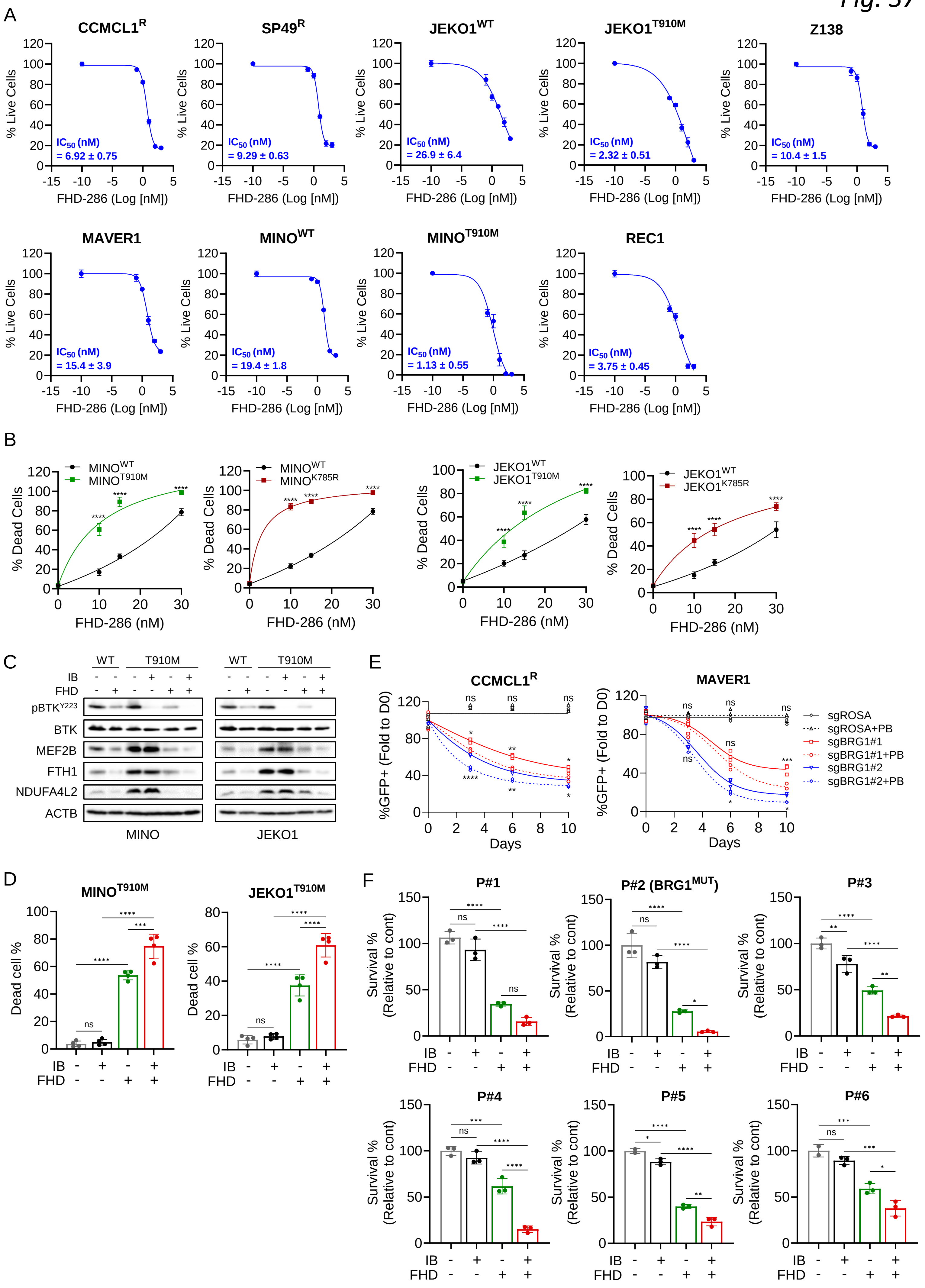
